# Novel mRNA-based VP8* vaccines against rotavirus are highly immunogenic in rodents

**DOI:** 10.1101/2023.03.29.534747

**Authors:** Sandro Roier, Vidya Mangala Prasad, Monica M. McNeal, Kelly K. Lee, Benjamin Petsch, Susanne Rauch

## Abstract

Despite the availability of live-attenuated oral vaccines, rotavirus remains a major cause of severe childhood diarrhea worldwide. Due to the growing demand for parenteral rotavirus vaccines, we developed novel mRNA-based vaccine candidates targeting the viral spike protein VP8*. Our monomeric P2 (universal T cell epitope)-VP8* mRNA design is equivalent to a protein vaccine currently in clinical development, while LS (lumazine synthase)-P2-VP8* was designed to form nanoparticles. Cyro-electron microscopy and western blotting-based data presented here suggest that proteins derived from LS-P2-VP8* mRNA are secreted *in vitro* and self-assemble into 60-mer nanoparticles displaying VP8*. mRNA encoded VP8* was immunogenic in rodents and introduced both humoral and cellular responses. LS-P2-VP8* induced superior humoral responses to P2-VP8* in guinea pigs, both as monovalent and trivalent vaccines, with encouraging responses detected against the most prevalent P genotypes. Overall, our data provide evidence that trivalent LS-P2-VP8* represents a promising mRNA-based next-generation rotavirus vaccine candidate.

## INTRODUCTION

Rotavirus, a non-enveloped segmented double-stranded RNA virus belonging to the *Reoviridae* family, is still the leading cause of severe dehydrating gastroenteritis in infants and children under 5 years of age worldwide. The vast majority of rotavirus-associated deaths, that amount to approx. 200.000 globally every year, occurs in low- and middle-income countries (LMICs)^1–6^. The currently available live-attenuated oral rotavirus vaccines demonstrated up to 97% protection against severe rotavirus gastroenteritis in clinical trials and have led to significant decline in the global burden of rotavirus-induced disease. However, in LMICs with high diarrheal mortality, lower efficacy in clinical trials and reduced real-world vaccine effectiveness were observed compared to high-income countries with low diarrheal mortality. The reason for the modest vaccine efficacy and effectiveness in LMICs is likely multifactorial and may also be related to the lower immunogenicity and efficacy of other oral vaccines demonstrated in these settings^4, 7–9^. With the additional, if low, risk of vaccination-associated intussusception reported for two live-attenuated oral rotavirus vaccines in post-marketing studies in high- and middle-income countries^10, 11^, there is growing interest in the development of non-replicating parenteral vaccines that might overcome the limitations of current rotavirus vaccines^12–14^. The most advanced candidate in this category is the protein vaccine P2- VP8*, which was developed at the NIH^15, 16^ and later advanced clinically by PATH^17–19^ to an ongoing phase 3 efficacy trial (ClinicalTrials.gov identifier: NCT04010448). This recombinant protein vaccine consists of the VP8* core domain fused to the universal CD4^+^ T cell epitope P2 derived from tetanus toxin to enhance immunogenicity. VP8* is a proteolytic cleavage product of the outer capsid spike protein VP4 that contains neutralizing epitopes and defines the rotavirus P genotype^15, 16, 19, 20^. Alum-adjuvanted monovalent and trivalent vaccine formulations of P2-VP8* containing the most clinically relevant VP8* genotypes P[8], P[6] and P[4]^1, 3, 14^ showed promising results in preclinical studies using mice^21^, guinea pigs^16, 22, 23^ and gnotobiotic pigs^16^. Since preclinical^16, 21, 22^ and clinical^17, 18^ findings demonstrated reduced neutralizing antibody responses to heterotypic rotaviruses, the trivalent P2-VP8* vaccine was selected for further clinical development. This trivalent formulation induced robust IgG and neutralizing responses to P[8], P[6] and P[4] rotaviruses in humans^19^.

In this work, we investigated the immunogenicity of two novel mRNA-based rotavirus vaccine candidates targeting VP8* in rodents. These injectable non-replicating vaccines were developed using CureVac’s proprietary RNActive^®^ technology^24–31^. The vaccine candidates consist of lipid nanoparticle (LNP)-formulated, chemically unmodified, and sequence-engineered mRNA. The vaccines include optimized non-coding regions that have previously been shown to greatly improve the immunogenicity and/or protective efficacy of an unmodified mRNA SARS-CoV-2 vaccine in mice^32^, rats^33^ and non-human primates^34^. The first mRNA vaccine candidate (P2-VP8*) encodes an equivalent design of the protein vaccine P2-VP8*, with differences in the underlying rotavirus P[8], P[6] and P[4] isolates. The second mRNA vaccine candidate (LS-P2-VP8*) encodes a multimerization domain derived from *Aquifex aeolicus* lumazine synthase (LS)^35^ fused to P2-VP8*. LS from this hyperthermophilic bacterium naturally self-assembles into 60-mer icosahedral nanoparticles^36^. LS-based approaches have successfully been employed in other vaccine studies to display antigens on the nanoparticle surface, thereby significantly improving the immunogenicity of the presented antigens^37–40^. The success of these studies is likely due to the fact that multimerization of antigens on a nanoparticle surface favors the generation of potent, long-lived immunoprotection in germinal centers through clustering of B cell receptors for multiple and simultaneous binding to the antigen epitopes^35, 41^.

Here we evaluated the immune responses induced by monovalent and trivalent formulations of the P2-VP8* and LS-P2-VP8* mRNA vaccines in mice and guinea pigs and compared them with those elicited by the corresponding alum-adjuvanted P2-VP8* protein vaccines. In addition, we characterized cell culture-derived nanoparticles translated from an mRNA encoding LS-P2-VP8* P[8] by high resolution cryo-electron microscopy (cryo-EM).

## RESULTS

### An mRNA vaccine encoding P2-VP8* P[8] is immunogenic in mice

In order to assess the immunogenicity of our mRNA-based rotavirus vaccine approach, we immunized BALB/c mice intramuscularly (IM) with either 6 µg of alum-adjuvanted P2-VP8* P[8] protein or 4 µg of LNP-formulated P2-VP8* P[8] mRNA vaccine. Mice were vaccinated twice (day 0 and day 21), and humoral and cellular immune responses were evaluated after one (day 21) or two vaccinations (day 42 and day 56).

Compared to sham-immunized animals, mice vaccinated with the protein vaccine showed significantly higher P2-VP8* P[8]-specific binding IgG1 and IgG2a antibody titers in serum after the second vaccination, with maximum geometric mean endpoint titers of 6.0 × 10^5^ for IgG1 on day 56 and 1.7 × 10^3^ for IgG2a on day 42 (Fig. 1a). In contrast, antigen-specific IgG1 and IgG2a titers were already significantly increased after the first immunization with the P2-VP8* P[8] mRNA vaccine. Binding antibody titers were boosted by a second immunization to peak geometric mean endpoint titers of 2.3 × 10^6^ (IgG1; day 56) and 6.0 × 10^5^ (IgG2a; day 42). A comparison between the protein and mRNA vaccines revealed significantly elevated IgG1 and IgG2a titers after one immunization as well as significantly increased IgG2a titers after the second immunization for the mRNA-based vaccine.

**Fig. 1.**
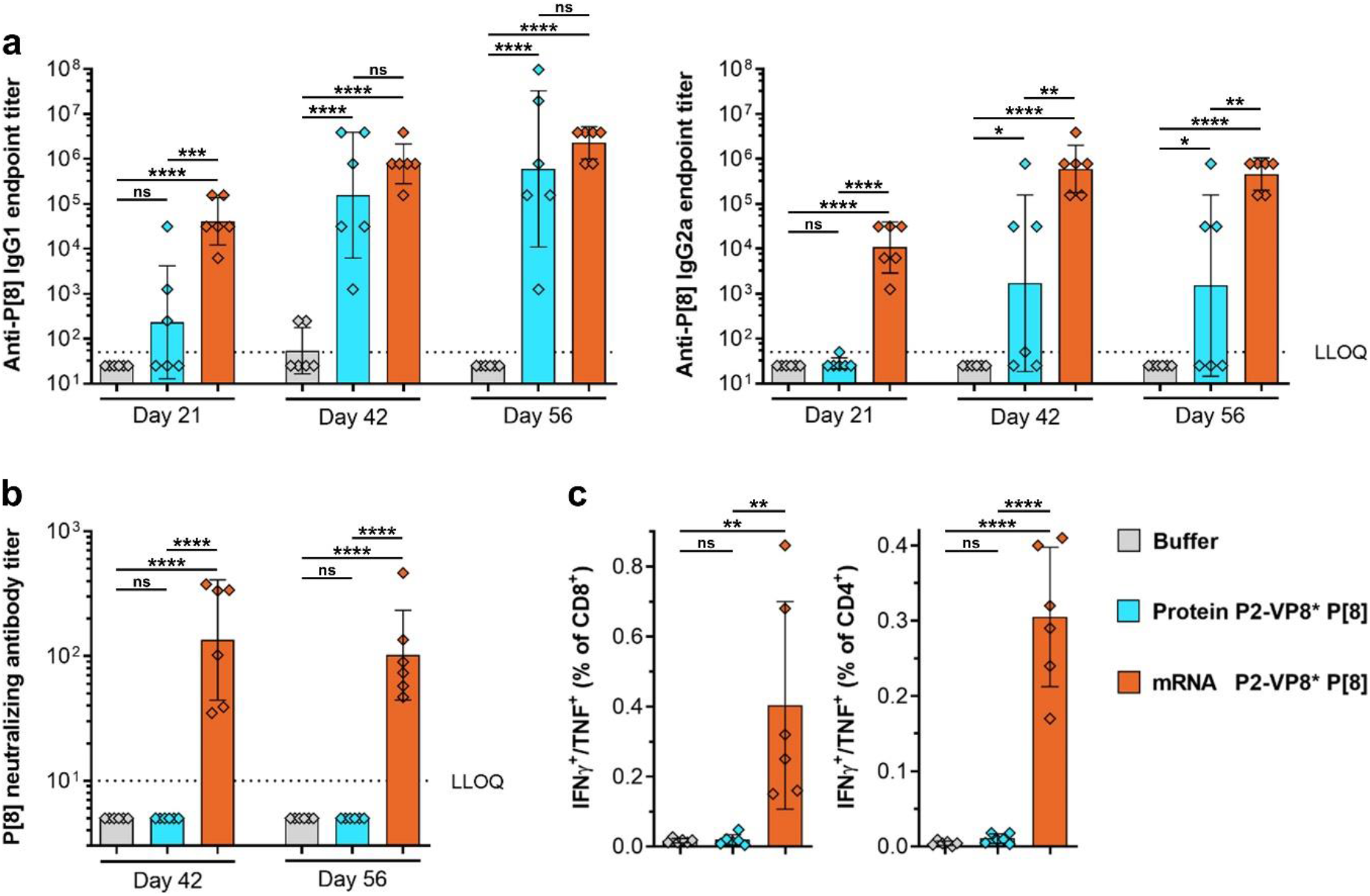
mRNA encoding P2-VP8* P[8] induced robust levels of both humoral and cellular immune responses in mice. Female BALB/c mice (n = 6/group) were vaccinated IM with 6 µg of alum-adjuvanted P2-VP8* P[8] protein or 4 µg of P2-VP8* P[8] mRNA vaccine on day 0 and day 21. Buffer (0.9% NaCl) vaccinated animals served as negative controls. Sera were collected on day 21, day 42 and day 56 and splenocytes were isolated on day 56. **a** P2-VP8* P[8]-specific binding antibodies are displayed as endpoint titers for IgG1 and IgG2a in serum upon one (day 21) or two vaccinations (day 42 and day 56). **b** P[8] genotype (Wa strain)-specific rotavirus-neutralizing antibody titers in serum upon two vaccinations (day 42 and day 56). **c** Multifunctional IFN-γ/TNF-positive CD4^+^ and CD8^+^ T cells were analyzed in splenocytes stimulated with VP8*-specific peptides followed by intracellular cytokine staining and detection by flow cytometry. Each square represents an individual animal and bars depict geometric means with geometric standard deviation (SD) in panel **a** and **b** and means with SD in panel **c**. Dotted lines indicate the lower limit of quantification (LLOQ). Values below the LLOQ (<50 for panel **a** and <10 for panel **b**) were set to half of the respective LLOQ. Individual readouts and time points were statistically analyzed using an ordinary one-way ANOVA followed by Tukey’s multiple comparison post test. Significant differences between groups are marked by asterisks (\**P*<0.05, ** *P*<0.01, *** *P*<0.001, **** *P*<0.0001; ns: not significant).

Induction of virus-neutralizing antibody (nAb) titers in serum was assessed three (day 42) or five weeks (day 56) after the second vaccination by measuring homotypic responses (P[8] genotype) to P[8] (Wa strain) rotavirus (Fig. 1b). For both time points tested, a significantly increased nAb response was detected after vaccination with the mRNA vaccine, with a maximum geometric mean neutralizing titer of 134 on day 42. All animals immunized with the protein vaccine showed no detectable nAb titers above the lower limit of quantification of 10.

To investigate the cellular immune response, splenocytes were isolated five weeks after the last immunization (day 56), stimulated with VP8*-specific peptides, and the induced multifunctional IFN-γ/TNF-positive CD4^+^ and CD8^+^ T cells were analyzed by intracellular cytokine staining and flow cytometry (Fig. 1c). Supplementary Fig. 1 illustrates the gating strategy used for T cell analysis. As observed with the nAb responses, only the mRNA vaccine yielded significant IFN-γ/TNF double-positive CD8^+^ and CD4^+^ T cell responses, with mean values of 0.4% and 0.3%, respectively. As expected^42^, the alum-adjuvanted protein vaccine induced only minor, non-significant T cell responses compared to the negative control group.

Overall, we demonstrated that mRNA encoding P2-VP8* P[8] induces robust levels of both humoral and cellular immune responses in mice.

### Proteins translated from mRNA encoding LS-P2-VP8* P[8] form nanoparticles in cell culture

To further improve the immunogenicity of mRNA encoded VP8*, we generated the fusion construct LS-P2-VP8*. This mRNA design features lumazine synthase, a 60-mer nanoparticle-forming protein from *Aquifex aeolicus*, fused to the N-terminus of P2-VP8* (Fig. 2a). First experiments evaluated the optimal length for a secreted form of VP8* in a pDNA-based screen (Supplementary Fig. 2). A VP8* construct featuring aa 41-223 was chosen as the basis for LS-P2-VP8*. This fragment was selected as the closest to the well characterized P2-VP8* design that demonstrated both solid VP8* expression (lysates) and secretion (supernatants) levels.

**Fig. 2.**
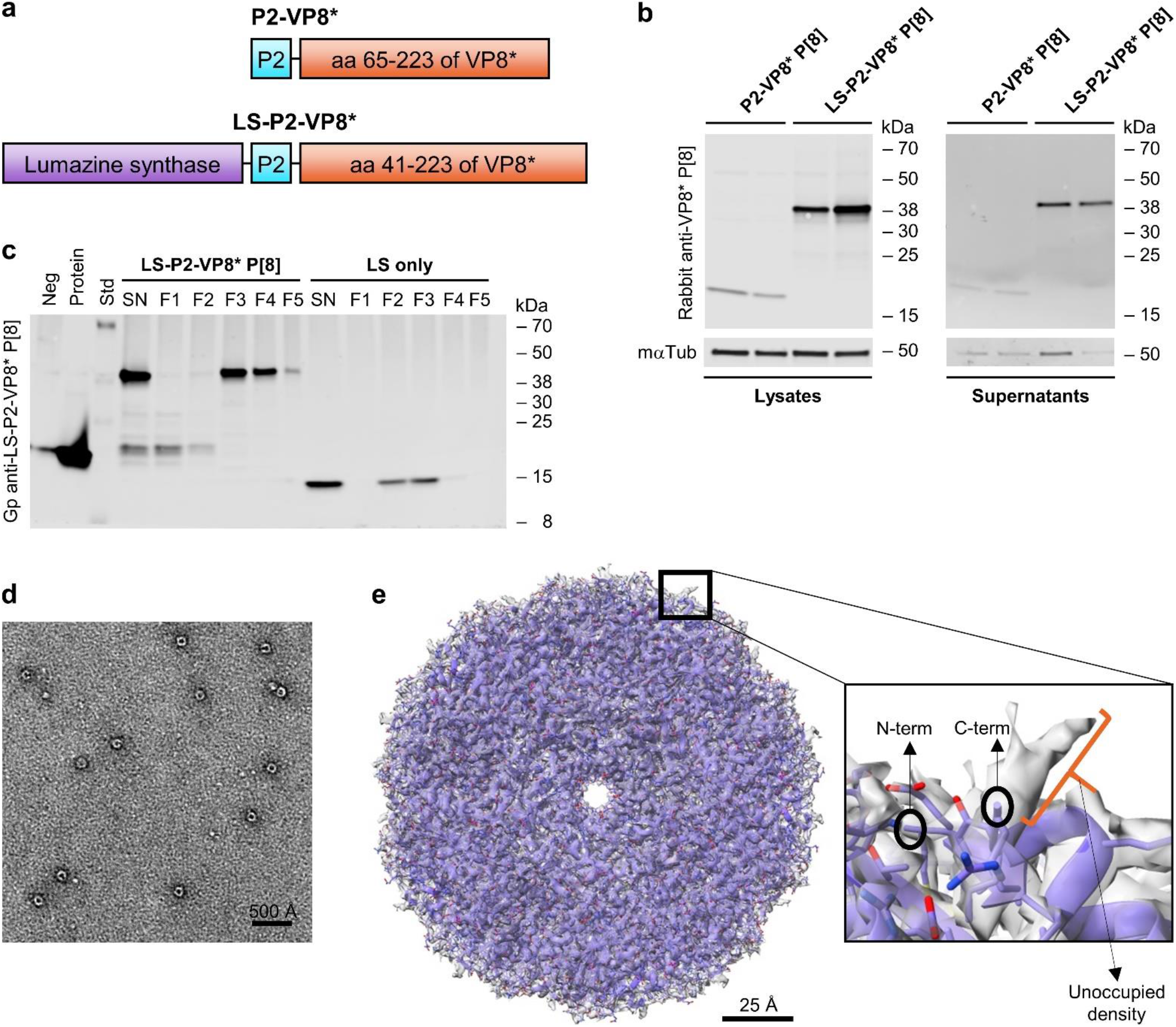
Proteins translated from an mRNA encoding LS-P2-VP8* P[8] were secreted and self-assembled into 60-mer nanoparticles. **a** Schematic drawings of mRNA encoded P2-VP8* and LS-P2-VP8* proteins. **b** HEK 293T cells were transfected with two independent mRNA batches each of P2-VP8* P[8] or LS-P2-VP8* P[8]. VP8* expression in cell lysates and supernatants were analyzed via western blotting 48 hours post transfection. Tubulin was employed as loading control. **c** Cell supernatants of HeLa cells transfected with LS-P2-VP8* P[8] or LS only mRNA were subjected to density gradient ultracentrifugation using an isopycnic iodixanol density gradient 48 hours post transfection. After ultracentrifugation, equivalent volumes of five density gradient fractions with increasing iodixanol concentrations (F1 to F5) or unpurified supernatants (SN) were analyzed for VP8* and/or LS expression via western blotting. A supernatant from HeLa cells transfected with water served as negative control (Neg) and 100 ng of P2-VP8* P[8] protein loaded to the gel was used as positive control (Protein). **d** Negative stain EM image of LS-P2-VP8* P[8] nanoparticles derived from HeLa cell culture supernatants. White is high density. Circular particles with hollow interior were observed. **e** Cryo-EM reconstruction of LS-P2-VP8* P[8] nanoparticles derived from HEK 293T cell culture supernatants with PDB model of LS fitted into it. Rectangular box on surface of nanoparticle is shown as zoomed inset. Both N-terminus and C- terminus of LS are located at the surface of the nanoparticle. P2-VP8* is not seen in the cryo-EM map, but a short unoccupied density is visible beyond the C-terminal residue of the LS structure, suggesting that P2- VP8* is not resolved in the cryo-EM density due to the flexible linker connecting P2-VP8* to LS. LS: lumazine synthase; aa: amino acid; mαTub: mouse anti-alpha tubulin antibody; Gp: guinea pig.

Two independent mRNA batches of P2-VP8* P[8] or LS-P2-VP8* P[8] were employed for testing the expression of mRNA encoded VP8* (Fig. 2b). Both mRNA constructs expressed proteins of the expected size of 20.4 kDa and 40.6 kDa, respectively. These experiments demonstrated enhanced *in vitro* protein expression in cell lysates and increased protein secretion into cell culture supernatants of LS-P2-VP8* P[8] compared to P2-VP8* P[8].

In order to characterize LS-P2-VP8* P[8] nanoparticles translated from mRNA in cell culture, supernatants of cells transfected with LS-P2-VP8* P[8] were purified by density gradient ultracentrifugation and analyzed by transmission electron microscopy (TEM). mRNA encoding lumazine synthase alone (LS only) was employed as control. Density gradient fractions with increasing iodixanol concentrations were analyzed for VP8* and/or LS expression using an LS-P2-VP8* P[8]-specific polyclonal antiserum (Fig. 2c) or exclusively for VP8* expression using a VP8* P[8]-specific polyclonal antibody (Supplementary Fig. 3). Both LS-P2-VP8* P[8] and LS only mRNA constructs yielded the expected monomeric protein sizes in the unpurified supernatants of 40.6 kDa and 16.7 kDa, respectively, when checked using SDS-PAGE and western blotting (Fig. 2c). Potential degradation products in the LS-P2-VP8* P[8] transfected supernatant were efficiently removed by density gradient ultracentrifugation, and the main band signal was found to accumulate in fractions 3 and 4, which corresponds to a mean iodixanol concentration of approximately 27% (19-36%). LS only-specific band signals could be detected with polyclonal antiserum generated upon vaccination with LS-P2-VP8* P[8] (compare Fig. 2c and Supplementary Fig. 3), and accumulated in density gradient fractions 2 and 3, corresponding to a mean iodixanol concentration of approximately 19% (11-27%).

To confirm the correct formation of LS-P2-VP8* P[8] nanoparticles, density gradient fractions 3 and 4 were pooled and subjected to TEM analysis. The pooled gradient fractions were analyzed using negative stain EM, which showed a uniform distribution of spherical particles with hollow interiors (Fig. 2d). Based on this, the samples were then vitrified for single particle cryo-EM analysis. A cryo-EM structure of LS-P2-VP8* P[8] nanoparticles was determined at 2.6Å resolution (at 0.143 F.S.C cutoff) (Fig. 2e and Supplementary Fig. 4). The atomic model of LS (PDB ID: 5MPP) was fitted into the cryo-EM map (Fig. 2e) without any modifications, confirming that the LS-P2-VP8* P[8] core nanoparticle structure agreed well with the previously published structure of a 60-mer LS nanoparticle^43^. In the 2D class averages generated from the LS-P2-VP8* P[8] nanoparticle dataset, image classes were observed that show diffuse, nonuniform density protruding from nanoparticle surface, which may correspond to the P2-VP8* portion (Supplementary Fig. 5). However, ordered density corresponding to P2-VP8* was not visible in the cryo-EM map (Fig. 2e). Only a short segment of unoccupied density in the cryo-EM map at the end of the C-terminus of LS was observed (Fig.2e) where the P2-VP8* moiety is expected to be present. Since LS and P2-VP8* are fused via a flexible protein linker, it seems likely that the P2-VP8* protein portion cannot be clearly visualized on the surface of the LS nanoparticle structure due to this flexibility.

Our data indicate that the protein translated from an mRNA encoding an LS nanoparticle domain fused to P2-VP8* P[8] is secreted *in vitro* and self-assembles into 60-mer nanoparticles.

### Monovalent VP8*-based mRNA vaccines induce long-lived immune responses in guinea pigs

Since the P2-VP8* P[8] protein vaccine elicited only modest immune responses in the mouse model and most preclinical studies with this vaccine were performed using guinea pigs^16, 22, 23^, this model system was chosen for further immunogenicity studies. Immune responses induced by 20 µg of P2-VP8* P[8] protein vaccine compared with 25 µg of an mRNA vaccine, i.e. P2-VP8* P[8] or LS-P2-VP8* P[8], were assessed in Dunkin Hartley guinea pigs upon two IM vaccinations on day 0 and day 21. Humoral immune responses were evaluated after one or two vaccinations and followed over a period of 6 months. Sham-immunized animals served as negative controls.

P2-VP8* P[8]-specific total serum IgG antibody titers peaked on day 42 in all vaccine groups and reached a stable plateau in all later measurements (Fig. 3a). A trend towards higher IgG titers after two vaccinations was seen in all groups including the buffer control group, possibly due to assay variations and increasing age of the animals leading to an elevated background signal. In contrast to the binding antibody responses observed in the mouse model, guinea pigs vaccinated with the protein vaccine showed higher IgG titers after two vaccinations on day 42 than the corresponding mRNA vaccine. The highest specific IgG titers were observed in animals immunized with the LS-P2-VP8* P[8] mRNA vaccine. This was already observed on day 21 after the first vaccination and consistently until the end of the observation period on day 196. Differences in specific IgG titers between vaccine groups remained constant over time.

**Fig. 3.**
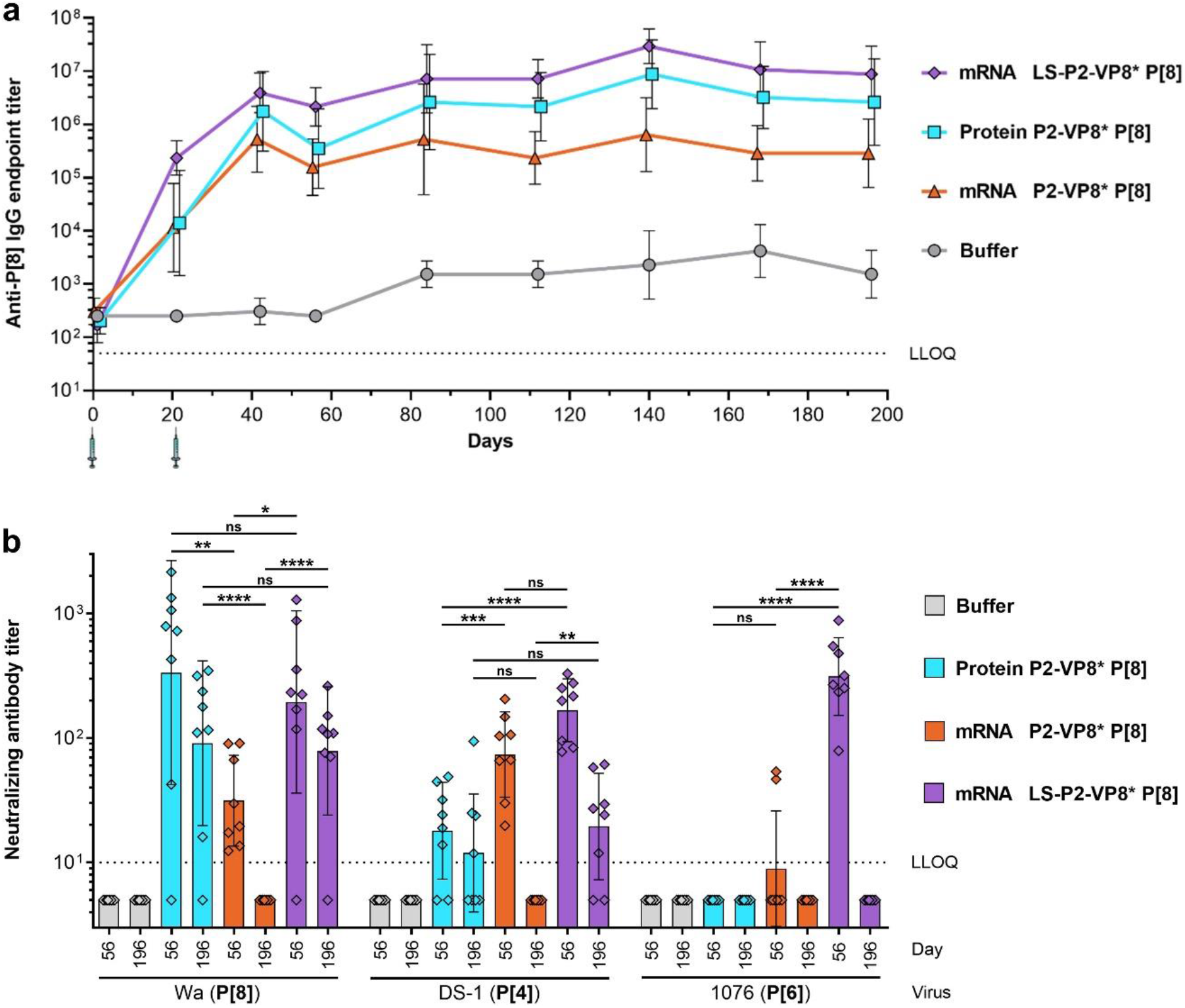
mRNA encoding LS-P2-VP8* P[8] induced high and long-lasting levels of humoral immunity in guinea pigs. Female Dunkin-Hartley guinea pigs (n = 8/group) were vaccinated IM with 20 µg of alum- adjuvanted P2-VP8* P[8] protein or 25 µg each of P2-VP8* P[8] or LS-P2-VP8* P[8] mRNA vaccines on day 0 and day 21. Buffer (0.9% NaCl) vaccinated animals served as negative controls. Sera were collected on different days as indicated. **a** P2-VP8* P[8]-specific binding antibodies are displayed as endpoint titers for total IgG in serum at the indicated time points. Each dot represents the geometric mean with geometric standard deviation (SD) of a group at a given time. Syringe symbols indicate the vaccination time points. **b** P[8] genotype (Wa strain), P[4] genotype (DS-1 strain) or P[6] genotype (1076 strain)-specific rotavirus- neutralizing antibody titers in serum upon two vaccinations (day 56 and day 196). Each square represents an individual animal and bars depict geometric means with geometric SD. Dotted lines indicate the lower limit of quantification (LLOQ). Values below the LLOQ (<50 for panel **a** and <10 for panel **b**) were set to half of the respective LLOQ. Individual readouts and time points in panel **b** were statistically analyzed using an ordinary one-way ANOVA followed by Šídák’s multiple comparison post test. Significant differences between groups are marked by asterisks (\**P*<0.05, ** *P*<0.01, *** *P*<0.001, **** *P*<0.0001; ns: not significant).

Induction of nAb titers in serum was assessed five weeks (day 56) or six months (day 196) after the second vaccination by measuring homotypic responses to P[8] genotype (Wa strain) and heterotypic responses to P[4] and P[6] genotypes (DS-1 and 1076 strains) (Fig. 3b). When comparing homotypic nAb responses, no significant differences were observed between groups immunized with protein vaccine and LS-P2-VP8* P[8] mRNA vaccine at either time point. Geometric mean neutralizing titers in both groups were higher on day 56 and decreased six months after the last vaccination from 335 (day 56) to 91 (day 196) and from 195 (day 56) to 79 (day 196) for the protein vaccine group and the LS-P2-VP8* P[8] mRNA vaccine group, respectively. In contrast to these groups, immunization with P2-VP8* P[8] mRNA yielded significantly lower geometric mean neutralizing titers of 31 on day 56, and no nAb titers above the lower limit of quantification were detectable on day 196.

Analysis of P[4]-specific heterotypic nAbs revealed overall higher titers in mRNA vaccine than in the protein vaccine groups. On day 56, significantly higher geometric mean titers of 167 and 74 were measured in the LS-P2-VP8* P[8] and P2-VP8* P[8] mRNA vaccine groups, respectively, compared with 18 in the protein vaccine group. On day 196, P[4]-specific nAb titers above the lower limit of quantification only remained detectable in animals vaccinated with the protein vaccine and the LS-P2-VP8* P[8] mRNA vaccine, with geometric mean titers of 12 and 19, respectively. P[6]-specific heterotypic nAb responses were detected only in the LS-P2-VP8* P[8] mRNA vaccine group on day 56, with a geometric mean neutralizing titer of 312.

Taken together, these data show that a monovalent mRNA vaccine encoding LS-P2-VP8* P[8] induces high and long-lasting binding antibody responses and is capable of eliciting nAbs not only against homotypic but also against heterotypic rotaviruses in guinea pigs. While the P2-VP8* P[8] protein vaccine performed comparably well in terms of homotypic immune responses, the LS-P2-VP8* P[8] mRNA vaccine was superior in inducing heterotypic nAb responses at the doses tested.

### Trivalent VP8*-based mRNA vaccines are immunogenic in guinea pigs

In order to develop a vaccine that is effective against the most prevalent disease-causing rotavirus genotypes P[8], P[4] and P[6]^1,3,14^, we next generated trivalent P2-VP8* or LS-P2-VP8* mRNA vaccines and tested their immunogenicity in guinea pigs. For this, P[6] or P[4] genotype-specific mRNAs corresponding to the described P[8] designs were produced and protein expression was confirmed in cell cultures (data not shown). Monovalent and trivalent mRNA vaccines were generated by encapsulating single mRNAs (genotype P[8], P[6] or P[4] of VP8*) or a pre-mix of all three mRNAs (1:1:1 weight ratio), respectively, in LNPs. These mRNA vaccines were compared to monovalent (P[8]) and trivalent (P[8], P[6] and P[4]; 1:1:1 weight ratio) alum-adjuvanted P2-VP8* protein vaccines. For this, Dunkin Hartley guinea pigs were immunized IM with 20 µg of the respective protein vaccine or 25 µg of the respective mRNA vaccine. For the trivalent LS-P2-VP8* vaccine, additional groups were vaccinated with a low dose of 3 µg (1 µg per mRNA). Animals were vaccinated three times (day 0, day 21, and day 42) or twice (day 21 and day 42), and humoral immune responses were evaluated in serum collected before vaccination on day 21 and day 42 or at the end of the study on day 70. Sham-immunized animals served as negative controls.

Overall, monovalent mRNA vaccines elicited the highest responses in homotypic levels of IgG antibodies, while heterotypic responses were comparatively lower (Fig. 4 and Supplementary Fig. 6 and 7). In contrast, the trivalent mRNA vaccines performed equally well across all three P genotype-specific IgG responses. P[8]-, P[6]-, and P[4]-specific antibody levels elicited by the mRNA vaccines were increased 12-fold on average by a third vaccination (compare Fig. 4c and Supplementary Fig. 6 and 7), whereas IgG titers induced by the trivalent P2-VP8* protein vaccine did not benefit from a third shot. 3 µg or 25 µg of trivalent LS-P2-VP8* mRNA vaccine performed similarly after one, two, or three vaccinations, concerning all P genotype-specific antibody responses. When comparing P[8]-specific IgG responses, no significant differences were found at any time point between groups immunized with the trivalent P2-VP8* protein vaccine and the trivalent LS-P2-VP8* mRNA vaccine (Fig. 4). Three administrations of the trivalent P2-VP8* mRNA vaccine were required to induce comparably high P[8]-specific IgG titers to the other two trivalent vaccines. Similar trends between trivalent protein and mRNA vaccine groups were observed for the P[6]-(Supplementary Fig. 6) and the P[4]-specific (Supplementary Fig. 7) IgG responses.

**Fig. 4.**
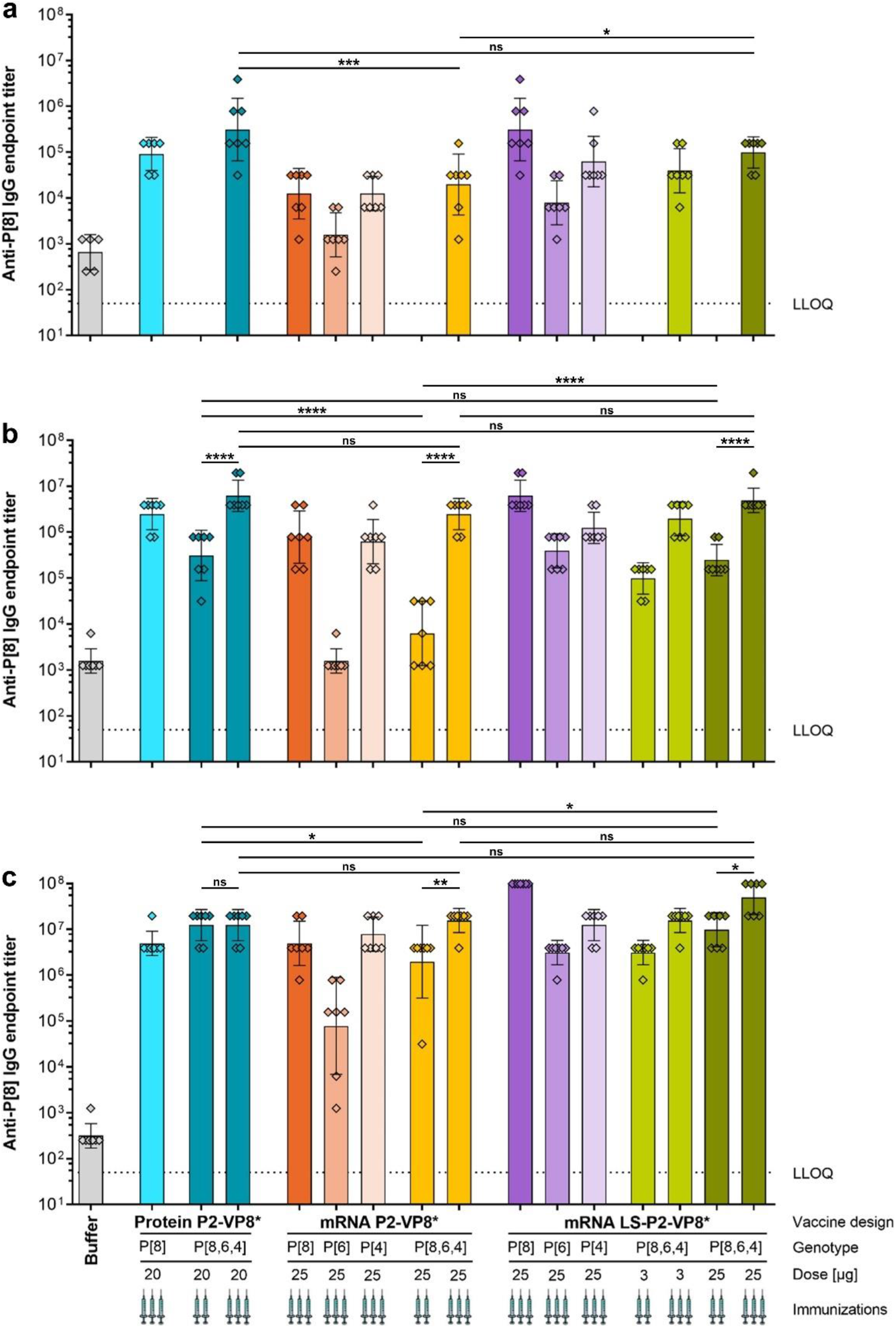
Trivalent LS-P2-VP8* mRNA vaccine encoding genotypes P[8], P[6] and P[4] of VP8* elicited high titers of binding antibodies against the P2-VP8* P[8] protein in guinea pigs. Female Dunkin- Hartley guinea pigs (n = 7/group) were vaccinated IM either three times on day 0, day 21, and day 42 or twice on day 21 and day 42 with different doses of monovalent and trivalent alum-adjuvanted P2-VP8* protein vaccines or monovalent and trivalent P2-VP8* or LS-P2-VP8* mRNA vaccines as indicated. The genotype(s) of the encoded VP8* employed for each group, the doses used as well as the number of immunizations administered (indicated by the number of syringe symbols) are displayed beneath each group. Trivalent vaccines are labeled P[8,6,4]. Buffer (0.9% NaCl) vaccinated animals served as negative controls. P2-VP8* P[8]-specific binding antibodies are displayed as endpoint titers for total IgG in serum collected on day 21 a, day 42 b, or day 70 c. Each square represents an individual animal and bars depict geometric means with geometric SD. Dotted lines indicate the lower limit of quantification (LLOQ). Individual time points were statistically analyzed using an ordinary one-way ANOVA followed by Šídák’s multiple comparison post test. Significant differences between groups are marked by asterisks (\**P*<0.05, ** *P*<0.01, *** *P*<0.001, **** *P*<0.0001; ns: not significant).

The nAb responses to P[8] (Wa strain), P[6] (1076 strain) or P[4] (DS-1 strain) rotaviruses were investigated in sera collected four weeks (day 70) after the second or third vaccination (Fig. 5). Overall, these results were in agreement with the binding antibody responses. The nAb levels elicited by the trivalent mRNA vaccines were significantly increased by a third vaccination, on average 19-fold for P2-VP8* and 3-fold for LS-P2-VP8*. However, nAb responses revealed more pronounced differences between monovalent and trivalent mRNA vaccines. For all P genotypes, the trivalent mRNA vaccines induced equal or higher nAb levels compared to the corresponding monovalent mRNA vaccine, despite employing only a third of the mRNA encoding for homotypic VP8*. The relatively low P[6]-specific nAb responses (Fig. 5b) induced by all monovalent mRNA vaccines underscore the need for a trivalent mRNA vaccine approach to support efficacious vaccination. There were no significant differences between the trivalent protein and LS-P2-VP8* mRNA vaccines. An exception was the P[6]-specific nAb responses after two vaccinations, where trivalent LS-P2-VP8* performed better. Groups vaccinated with either 3 µg or 25 µg per dose of trivalent LS-P2-VP8* vaccine performed overall comparably. Among trivalent protein and mRNA vaccine groups, the LS-P2-VP8* vaccinated groups consistently exhibited the highest geometric mean neutralizing titers. A third vaccination was necessary for the trivalent P2-VP8* mRNA vaccine in order to match at least the P[8]-(Fig. 5a) and P[4]-specific (Fig. 5c) nAb titers elicited by the other two trivalent vaccines.

**Fig. 5.**
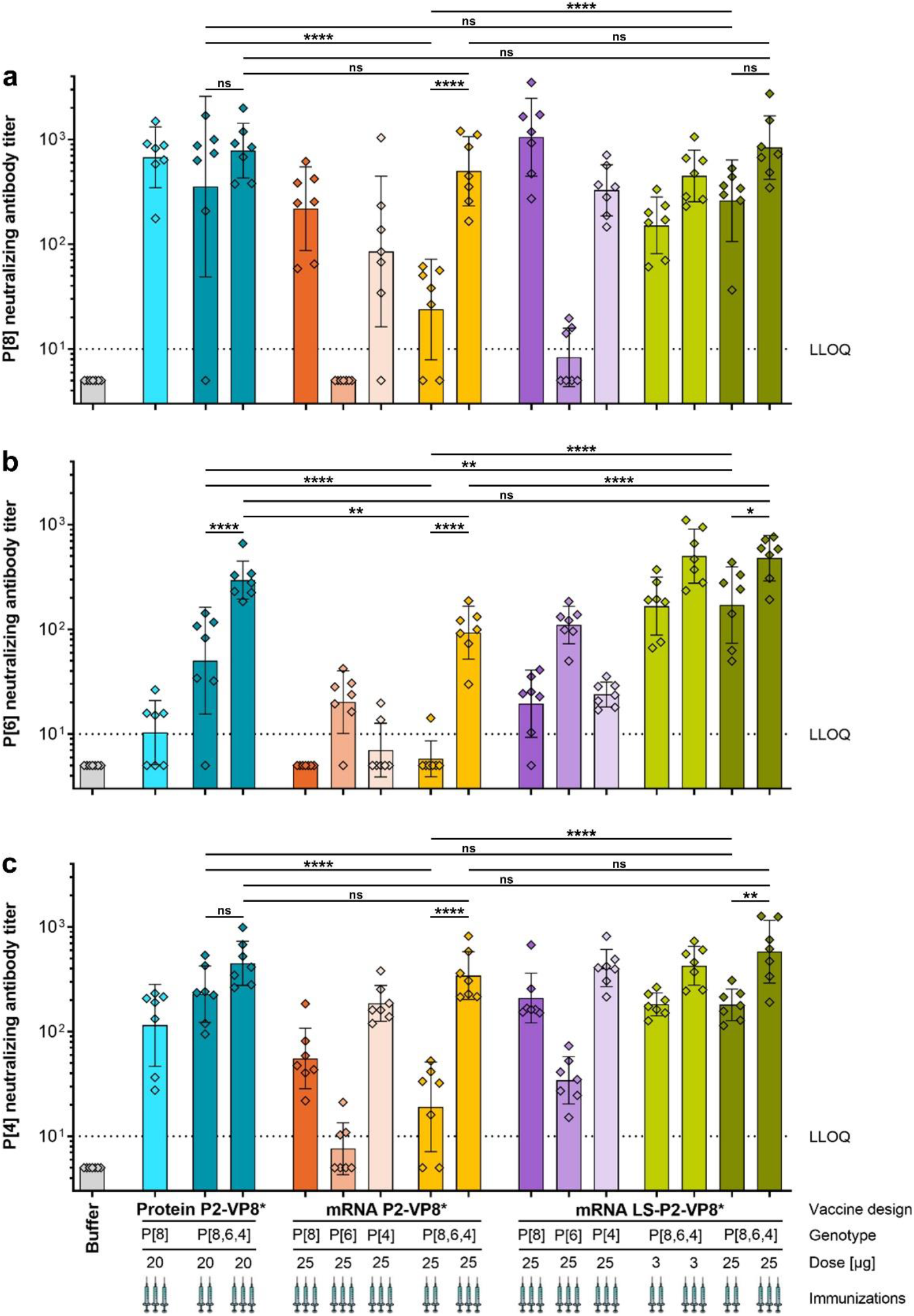
Trivalent LS-P2-VP8* mRNA vaccine encoding genotypes P[8], P[6] and P[4] of VP8* elicited high titers of functional antibodies against P[8], P[6] and P[4] rotavirus genotypes in guinea pigs. Female Dunkin-Hartley guinea pigs (n = 7/group) were vaccinated IM either three times on day 0, day 21, and day 42 or twice on day 21 and day 42 with different doses of monovalent and trivalent alum-adjuvanted P2-VP8* protein vaccines or monovalent and trivalent P2-VP8* or LS-P2-VP8* mRNA vaccines as indicated. The genotype(s) of the encoded VP8* employed for each group, the doses used as well as the number of immunizations administered (indicated by the number of syringe symbols) are displayed beneath each group. Trivalent vaccines are labeled P[8,6,4]. Buffer (0.9% NaCl) vaccinated animals served as negative controls. Sera were collected on day 70. a P[8] genotype (Wa strain), b P[6] genotype (1076 strain) or c P[4] genotype (DS-1 strain)-specific rotavirus-neutralizing antibody titers in serum. Each square represents an individual animal and bars depict geometric means with geometric SD. Dotted lines indicate the lower limit of quantification (LLOQ). Values below the LLOQ (<10) were set to half of the LLOQ. Individual readouts were statistically analyzed using an ordinary one-way ANOVA followed by Šídák’s multiple comparison post test. Significant differences between groups are marked by asterisks (\**P*<0.05, ** *P*<0.01, **** *P*<0.0001; ns: not significant).

In summary, we showed that mRNA vaccines induce nAb responses in guinea pigs against all P genotypes tested. nAb levels elicited by monovalent or trivalent LS-P2-VP8* mRNA vaccines were equivalent or higher than those induced by P2-VP8* protein vaccines. Our results point to an advantage of trivalent over monovalent VP8*-based rotavirus mRNA vaccines.

## DISCUSSION

The present study is the first to report mRNA-based rotavirus vaccine candidates and shows promising immunogenicity results in rodent models. The viral surface protein VP8*, responsible for viral attachment to the host cell^1^, was employed as a basis for mRNA vaccine development. In contrast to the majority of previously described mRNA vaccines that are mostly directed against enveloped viruses^27, 28, 30, 31, 44–46^, the encoded antigen VP8* is not a trimeric glycoprotein on the cell surface but is derived from a non-enveloped virus and remains a cytoplasmic monomer when expressed in the absence of further viral proteins^1^. These characteristics render VP8* a suboptimal target for humoral responses upon expression via mRNA. In addition to an mRNA vaccine that expresses wildtype VP8* fused to the universal T cell epitope P2, we therefore developed a vaccine design aimed to achieve improved antigen characteristics, i.e., the secreted nanoparticle-forming vaccine LS-P2-VP8*.

Indeed, our data demonstrated superior immunogenicity of LS-P2-VP8* compared to mRNA encoded monomeric VP8* in rodents. mRNA encoded LS-P2-VP8* induced consistently higher antibody levels than P2-VP8* mRNAs in guinea pigs, which held true over a 6-month period and for both mono-and multivalent vaccines. The LS nanoparticle-forming domain encoded in LS-P2-VP8* had previously been employed in the context of RNA replicon vaccines encoding HIV antigens^38^ and protein-based approaches have used norovirus-derived nanoparticles to display multimeric VP8*^47–49^. Our new LS-based VP8* design aims to support the formation of 60-mer nanoparticles that mimic the natural display of VP8* on the viral surface. Data presented here strongly suggest that proteins derived from mRNA encoded LS-P2-VP8* are secreted *in vitro* and self-assemble into 60-mer nanoparticles with VP8* displayed on their surface. This conclusion is based on (1) secretion of monomeric LS-P2-VP8* proteins of the expected molecular weight under denaturing conditions, (2) a different density gradient behavior of LS-P2-VP8* encoded proteins compared to proteins derived from a construct encoding LS alone, (3) the confirmation by cryo-EM that the LS-P2-VP8* core nanoparticle structure is consistent with the published 60-mer structure of the LS nanoparticle^43^, and (4) high virus-neutralizing as well as P2-VP8*-specific binding antibody responses induced by LS-P2-VP8* mRNA vaccines in guinea pigs. Of note, although we were able to visualize localized halos of density consistent with the expected P2-VP8* display on the nanoparticle surface in 2D class averages of the cryo-EM data (Supplementary Fig. 5), we were unable to clearly delineate the P2-VP8* part on the cryo-EM structure of the nanoparticle. The fusion of P2-VP8* to LS via a flexible linker likely prevented analysis of VP8* at high resolution.

The multimeric presentation of antigens in well-ordered arrays with defined orientations on a nanoparticle surface can resemble viral particles in size, shape, and clustering of the presented antigen. Such nanoparticle vaccine designs may provide enhanced antigen stability and/or immunogenicity^13^. Our LS-P2-VP8* design likely displays 60 P2-VP8* protrusions on its icosahedral nanoparticle surface. This resembles an icosahedral rotavirus virion that also features 60 protrusions in the form of spikes on its surface^50, 51^. However, instead of only one VP8* protein present at the tip of each protrusion of the LS-P2-VP8* nanoparticle, the rotavirus virion harbors two VP8* fragments located at the tip of each spike. The 60-mer LS-P2-VP8* nanoparticle core is ∼15 nm in diameter with ∼2 nm distance between adjacent VP8* protein as measured between the positions of C-terminal LS residue from the cryo-EM structure. The rotavirus virion is approximately 100 nm in diameter^52^ and the VP8* molecules between individual spikes are ∼18 nm apart. Thus VP8* on the LS-based particles is more densely arrayed and closely spaced than on the native virion. It is unlikely that the two Fab arms of an IgG or a B cell receptor, with an optimal span of ∼13 nm^53^ can readily span the distance between two inter-spike epitope sites on a rotavirus particle, and the 3-4 nm intra-spike VP8* positioning may be sterically incompatible with two Fabs from a single IgG binding. By contrast some subset of VP8* copies on LS-P2-VP8* may be compatible with bivalent antibody or B cell receptor binding, potentially contributing to the high immunogenicity of the LS-P2-VP8* mRNA vaccines.

As observed by others^35, 40, 54, 55^ we found that LS-P2-VP8* vaccinated animals induce antibodies against the LS nanoparticle scaffold (compare Fig. 2c and Supplementary Fig. 3). It is currently unclear whether such anti-LS immune responses negatively impact the efficacy of LS-based vaccines by diminishing antigen-specific immune responses upon repeated vaccination. However, in some previous studies that found anti-carrier antibodies to LS or bacterial ferritin nanoparticles, no adverse effects on vaccine immunogenicity were observed^40, 54–57^. An ongoing phase 1 clinical trial (NCT05001373) evaluating the safety and immunogenicity of LS-based HIV mRNA vaccines is likely to shed more light on the LS-specific immune responses elicited in humans after repeated vaccinations.

To assess if our mRNA-based vaccines can profit from a multivalent approach, we tested monovalent P[8], P[6] and P[4] VP8* designs compared to the corresponding trivalent vaccines. For monovalent vaccines, different degrees of heterotypic immune responses were detected depending on the P genotype. Comparatively strong heterotypic P[4] and P[8] responses were induced by P[8]-and P[4]-specific mRNA vaccines, respectively, in agreement with the relatively small evolutionary distance between these genotypes. Lower levels of cross-reactivity was seen for P[6], which is genetically more distant to P[4] and P[8] genotypes^49, 58^. Homotypic nAb responses to monovalent P[6] mRNAs remained comparatively low. It is conceivable that nAb induction was affected by sequence differences between P[6] VP8* encoded in LS-P2-VP8* and VP8* present in the virus employed for the neutralization assay, which differ in 13 of 183 amino acids. Vaccination with multivalent vaccines demonstrated overall improved immunogenicity: both trivalent mRNA vaccines elicited equal or higher immune responses than the respective homotypic monovalent mRNA vaccines, despite containing a third of the mRNA encoding for homotypic VP8*. The effect was most pronounced for nAb responses against P[6] that benefited most from a combination of multivalent mRNA vaccines and nanoparticle-forming design. Taken together, our data support a trivalent VP8*-based mRNA vaccine approach to broaden efficacy against the most prevalent disease-causing rotavirus genotypes P[8], P[4] and P[6]^1, 3, 14^.

There are currently no correlates of protection established for live-attenuated oral rotavirus vaccines, let alone for non-replicating parenteral vaccines. It seems likely that these vaccine types have different protective mechanisms and thus feature distinct correlates of protection^59^. Therefore, it is unclear how the promising preclinical results of mRNA-based rotavirus vaccines will translate to humans. Unfortunately, the ongoing phase 3 clinical trial (NCT04010448) assessing safety and efficacy of the trivalent P2-VP8* protein vaccine met futility criteria at interim analysis as the protein vaccine demonstrated no superior efficacy to a licensed oral rotavirus vaccine^59^. However, given the fundamental differences in immune responses elicited by protein and mRNA vaccines^60–62^, it seems conceivable that an mRNA vaccine could perform differently in humans despite the shared antigen. Indeed, mRNA encoded VP8* was able to induce both VP8*-specific T cell responses in mice that were absent in protein vaccinated groups and high levels of heterotypic nAbs in guinea pigs. The induction of both humoral and cellular immune responses may be important to increase protective efficacy against rotavirus disease^1, 3, 63^. Comparison of humoral responses induced by mRNA and protein vaccines revealed highly competitive responses induced by mRNA vaccines in guinea pigs: a 3 µg dose of trivalent LS-P2-VP8* mRNA elicited comparable responses to 20 µg of trivalent P2-VP8* protein. nAb levels induced by the trivalent protein vaccine in our study are in good agreement with those observed in a recently published guinea pig study after three vaccinations^23^. Furthermore, data generated in a gnotobiotic pig model of human rotavirus infection and diarrhea demonstrated that vaccination with trivalent LS-P2-VP8* mRNA reduced viral shedding after challenge to a higher degree than trivalent P2-VP8* protein vaccine (Hensley, C. *et al*.; manuscript in preparation).

To our knowledge, our data are the first to demonstrate immunogenicity of an mRNA vaccine against a non-enveloped virus and support further development of trivalent LS-P2-VP8* as a promising mRNA-based vaccine against rotavirus.

## METHODS

### mRNA vaccines

The mRNA vaccines are based on CureVac’s RNActive^®^ platform and do not contain chemically modified nucleosides. They comprise a 5′ cap1 structure, a 5′ untranslated region (UTR) from the human hydroxysteroid 17-beta dehydrogenase 4 gene (*HSD17B4*), a GC-enriched open reading frame, a 3′ UTR from the human proteasome 20S subunit beta 3 gene (*PSMB3*), a histone stem-loop (does not apply to the mRNA vaccine used in the mouse study), and a poly(A) tail. LNP encapsulation of mRNA was performed using LNP technology from Acuitas Therapeutics (Vancouver, Canada). LNPs are composed of an ionizable amino lipid, phospholipid, cholesterol, and a PEGylated lipid. The P2-VP8* mRNA vaccines encode a CD4^+^ T cell epitope (P2) of the tetanus toxin^20^ (aa residues 830-844; NCBI reference sequence: WP_011100836.1) N-terminally linked to a P genotype-specific truncated rotavirus VP8* subunit protein (aa 65-223 of VP4) by a GSGSG linker. The LS-P2-VP8* mRNA vaccines encode full-length lumazine synthase (NCBI reference sequence: WP_010880027.1) from *Aquifex aeolicus* containing two point mutations (C37A to remove an unpaired cysteine and N102D to remove a glycosylation site^38, 64^) N-terminally linked via (GGS)_4_-GGG to P2-VP8*, which is designed as described above, except that a longer version of VP8* (aa 41-223 of VP4) is encoded. P genotype-specific VP8* sequences derive from the following human rotavirus A strains: BE1128 for P[8] VP4 genotype (GenBank accession number: JN849135.1), BE1322/F01322 for P[6] VP4 genotype (GenBank accession number: JF460826.1), and BE1058 for P[4] VP4 genotype (GenBank accession number: JN849123.1). Trivalent mRNA vaccines encoding genotypes P[8], P[6] and P[4] of VP8* were formulated by mixing the respective mRNAs in a 1:1:1 weight ratio prior to LNP encapsulation.

### Plasmid DNAs

pDNAs used to study the impact of VP8* length on protein expression and secretion were generated by cloning DNA sequences encoding the human tissue plasminogen activator signal peptide (tPA-SP; aa 1-21 of tPA; GenBank accession number: CAA00642.1) in combination with different amino acid lengths of VP8* P[8] into mammalian expression vector pcDNA3.1(+).

### Protein vaccines

The recombinant fusion proteins P2-VP8* P[8], P2-VP8* P[6], and P2-VP8* P[4] expressed in *Escherichia coli* were kindly provided by PATH (Seattle, WA, USA) and consist of tetanus toxin T cell epitope P2 linked to aa 65-223 (for P[8]) or aa 64-223 (for P[6] and P[4]) of VP4 from rotavirus strain Wa (P[8]), 1076 (P[6]), or DS-1 (P[4]), respectively, using a GSGSG linker as previously described^15–17, 19, 22, 65^. The protein vaccines were alum-adjuvanted by adsorption of the recombinant fusion proteins to aluminum hydroxide (Alhydrogel^®^; Croda, Frederikssund, Denmark) as previously described^22, 66^. Briefly, two-fold working stocks containing 120 µg/mL of either monovalent P2-VP8* P[8] or trivalent P2-VP8* antigens (1:1:1 weight ratio; 40 µg/mL per antigen) in 0.5 mM PO_4_ in 0.9% saline were combined with an equal volume of an Alhydrogel^®^ suspension (2.24 mg Al/mL diluted in 0.5 mM PO_4_ in 0.9% saline) using gentle mixing for 20-24 hours to achieve formulations with 60 µg/mL total monovalent or trivalent P2-VP8* antigen containing aluminum hydroxide at 1.12 mg Al/mL in 0.5 mM phosphate buffer in 0.9% saline (pH 6.8).

### Animal studies

Female BALB/cAnNRj mice (Janvier Labs, France) were provided and handled by Preclinics Gesellschaft für präklinische Forschung mbH (Potsdam, Germany). All in-life experimental procedures undertaken during the course of the mouse immunization study were conducted in accordance with German laws and guidelines for animal protection and appropriate local and national approvals as well as in accordance with Directive 2010/63/EU of the European Parliament and of the Council of 22 September 2010 on the protection of animals used for scientific purposes. Animals were acclimated for at least 1 week before any procedures were carried out and were 6-8 weeks old at the start of the study. Mice (n = 6/group) were injected intramuscularly (IM) with 6 µg (112 µg Al) of alum-adjuvanted P2-VP8* P[8] protein (100 µL; 25 µL each into the left and right *M. tibialis* and *M. gastrocnemius*) or with 4 µg of P2-VP8* P[8] mRNA vaccine (25 µL into the left or right *M. tibialis*) on day 0 and day 21. Buffer vaccinated animals (25 µL of 0.9% NaCl into left or right *M. tibialis*) served as negative controls. Blood samples were collected into Z-clot activator microtubes (Sarstedt) by retro-orbital bleeding on day 21, day 42 and day 56. The samples were incubated for 0.5-1 hour at room temperature (RT), and then sera were obtained by collecting the supernatant after centrifugation (5 min; 10,000 × g; RT). At the end of the study on day 56, splenocytes were isolated by mechanical disruption of the spleen using 40 µm cell strainers. Red blood cells were lysed and splenocytes were washed, resuspended in supplemented medium and finally stored as single-cell suspensions at −80°C until use.

Female Dunkin Hartley guinea pigs (Envigo RMS, The Netherlands) were provided and handled by Covance Laboratories Ltd. (now Labcorp, Huntingdon, UK). All in-life experimental procedures undertaken during the course of the guinea pig immunization studies were subject to the provisions of the United Kingdom Animals (Scientific Procedures) Act 1986 Amendment Regulations 2012 (the Act). Animals were acclimated for at least 1 week before any procedures were carried out and were 6-7 weeks old at the start of the first study and 10-11 weeks old at the start of the second study. For the first study testing the longevity of immune responses induced by monovalent VP8*-based mRNA vaccines, guinea pigs (n = 8/group) were injected IM with 20 µg (374 µg Al) of alum-adjuvanted P2-VP8* P[8] protein (334 µL; 167 µL each into the left and right *M. gastrocnemius*) or with 25 µg each of P2-VP8* P[8] or LS-P2-VP8* P[8] mRNA vaccines (100 µL into the left or right *M. gastrocnemius*) on day 0 and day 21. Buffer vaccinated animals (100 µL of 0.9% NaCl into left or right *M. gastrocnemius*) served as negative controls. Blood samples were collected on day 1, day 21, day 42, day 56, day 84, day 112, day 140, day 168, and day 196 (end of study). For the second study testing the immunogenicity of trivalent VP8*-based mRNA vaccines, guinea pigs (n = 7/group) were injected IM either three times on day 0, day 21, and day 42 or twice on day 21 and day 42 with 20 µg (374 µg Al) each of monovalent (P[8]) or trivalent alum-adjuvanted P2-VP8* protein vaccines (334 µL; 167 µL each into the left and right *M. gastrocnemius*) or with 3 µg or 25 µg each of monovalent or trivalent P2-VP8* or LS-P2-VP8* mRNA vaccines (100 µL into the left or right *M. gastrocnemius*). Buffer vaccinated animals (100 µL of 0.9% NaCl into left or right *M. gastrocnemius* on day 0, day 21, and day 42) served as negative controls. Blood samples were collected on day 21, day 42, and day 70 (end of study). For both guinea pig studies, blood samples were collected via the saphenous vein or via cardiac puncture (end point only). The samples were left to clot for 1-2 hours at RT, and then sera were obtained by collecting the supernatant after centrifugation (10 min; 2,000 × g; 4°C).

The protein vaccine doses used in the respective animal model were derived from previous mouse and guinea pig studies performed by others^15, 21, 23^. The dosages used for the mRNA vaccines were estimated based on accumulated learnings from previous projects at CureVac. Of note, equal doses of protein and mRNA vaccine injected are not expected to result in equivalent amounts of antigen presented *in vivo*, since mRNA-based expression is highly dependent on the cell type-specific quality of mRNA vaccine uptake, antigen expression and/or secretion^67, 68^.

### Quantification of binding antibodies

P2-VP8* P[8]-, P2-VP8* P[6]-, or P2-VP8* P[4]-specific binding antibodies in sera of vaccinated mice (IgG1 and IgG2a) or guinea pigs (total IgG) were assessed by indirect enzyme-linked immunosorbent assay (ELISA). 96-well MaxiSorp ELISA plates (black) were coated with 0.5 µg/mL of a recombinant P genotype-specific P2-VP8* protein for 4-5 hours at 37°C. Plates were washed and blocked overnight at 4°C with 1% milk in PBS/0.05% Tween-20. Serum samples were added in serial dilution and incubated for 2-4 hours at RT. After washing, plates were incubated with either biotinylated rat anti-mouse IgG1 antibody (1:300; BD Biosciences, Cat. 550331), biotinylated rat anti-mouse IgG2a antibody (1:300; BD Biosciences, Cat. 550332), or horseradish peroxidase (HRP)-conjugated rabbit anti-guinea pig IgG (H+L) antibody (1:500; Invitrogen, Cat. PA1-28597) in blocking buffer for 1-1.5 hours at RT. For detection of biotinylated antibodies, plates were washed and HRP-conjugated streptavidin (1:1,000; BD Biosciences, Cat. 554066) in blocking buffer was added for 30 min at RT. Finally, plates were washed, Amplex^TM^ UltraRed reagent (1:200; Invitrogen, Cat. A36006) with H_2_O_2_ (1:2,000) was added, and fluorescence was detected after 45-60 min using a BioTek SynergyHTX plate reader (excitation 530/25, emission detection 590/35, sensitivity 45). The ELISA endpoint titers were defined as the highest reciprocal serum dilution that yielded a signal above the mean background signal plus 3-fold SD.

### Serum rotavirus neutralization assay

The rotavirus strains Wa (G1P[8]), 1076 (G2P[6]) and DS-1 (G2P[4]) were used in this assay. These viruses were originally obtained from the National Institutes of Health and a stock was grown for use in the assay. The rotavirus serum neutralizing antibody assay was performed by titrating a respective serum sample and incubating it with a constant amount of a specific strain of rotavirus to allow neutralization of the virus. The rotavirus/serum mix was then incubated on a rotavirus susceptible cell line, MA104 (monkey kidney) cells, cultured in 96-well plates for 1 hour to allow adsorption of the virus. The plates were washed to remove serum and then overlayed with media. Control wells on each 96-well plate received either diluted virus without sera (virus control) or no virus or sera (cell controls). After an overnight incubation allowing for rotavirus growth, the cells were frozen, thawed to lyse them, and then resuspended. The relative amount of unneutralized rotavirus antigen was determined using an antigen capture format ELISA to detect rotavirus. The ELISA plates were coated with rabbit anti-rotavirus IgG antibody. The resuspended lysates from the neutralization part of the assay were added and incubated. Following a blocking step with nonfat milk, guinea pig anti-rotavirus antiserum was added to the wells to detect captured rotavirus. Horseradish peroxidase (HRP) conjugated rabbit anti-guinea pig IgG (Jackson ImmunoResearch, West Grove, PA, USA) was added to detect the guinea pig anti-rotavirus antibody. The wells were developed with o-Phenylenediamine (SigmaAldrich), a peroxidase sensitive colorimetric substrate. Optical density of each well was read as absorbance at 492 nm. The amount of rotavirus present in the resuspended lysate from each well is inversely related to the amount of neutralizing antibody present in the serum. Each serum dilution series was modeled using a logistic regression function. For each fitted curve the reciprocal serum dilution which corresponds to a 40% response (ED_40_), compared to the virus controls, was determined and reported as the titer. The ED_40_ represents the titer of the serum against a given virus, which represents a 60% reduction in amount of virus^69^.

### T cell analysis

The induction of antigen-specific T cells was determined using intracellular cytokine staining in combination with flow cytometry. For this purpose, mouse splenocytes were thawed and 2 × 10^6^ cells per well (200 µL) were stimulated for 5-7 hours at 37°C using the following VP8*-specific peptides at 5 µg/mL each: FYIIPRSQE, KYGGRVWTF, VYESTNNSD, FYNSVWTFH, GFMKFYNSV, SDFWTAVIAVEPHVN, TNKTDIWVALLLVEP, and HKRTLTSDTKLAGFM. After 1 h, GolgiPlug^TM^ (BD Biosciences, Cat. 555029) was added in a dilution of 1:200 (50 µL) to the splenocytes to inhibit the secretion of cytokines. After stimulation, splenocytes were centrifuged, resuspended in supplemented medium, and stored overnight at 4°C. The next day splenocytes were washed twice in PBS and stained with LIVE/DEAD^TM^ fixable aqua dead cell stain kit (Invitrogen, Cat. L34957) for 30 min at 4°C. After an additional washing step in PBS with 0.5% BSA, cells were surface stained for Thy1.2 (FITC rat anti-mouse CD90.2 (Thy1.2); 1:200; BioLegend, Cat. 140304), CD4 (V450 rat anti-mouse CD4; 1:200; BD Biosciences, Cat. 560468) and CD8 (APC-H7 rat anti-mouse CD8a; 1:100; BD Biosciences, Cat. 560182) and incubated with FcγR-block (rat anti-mouse CD16/CD32; 1:100; Invitrogen, Cat. 14-0161-85) in PBS with 0.5% BSA for 30 min at 4°C. Subsequently, splenocytes were washed and fixed using Cytofix/Cytoperm^TM^ solution (BD Biosciences, Cat. 554722) for 20 min at RT in the dark. After fixation, cells were washed in perm buffer (PBS, 0.5% BSA, 0.1% Saponin) and stained for IFN-γ (APC rat anti- mouse IFN-γ; 1:100; BD Biosciences, Cat. 554413) and TNF (PE rat anti-mouse TNF alpha, 1:100; Invitrogen, Cat. 12-7321-82) for 30 min at RT. After intracellular cytokine staining, splenocytes were washed in perm buffer and resuspended in PFEA buffer (PBS, 2% FCS, 2 mM EDTA, 0.01% sodium azide). Finally, splenocytes were analyzed by flow cytometry on a Canto II flow cytometer (BD Biosciences) and the generated data were analyzed using FlowJo software (Tree Star, Inc.; Ashland, OR, USA).

### Generation of VP8*-specific polyclonal antibodies

Genotype-specific polyclonal antibodies against VP8* were generated by Aldevron Freiburg GmbH (now Genovac, Freiburg, Germany) by immunizing two rabbits each with either a VP8* P[8]/VP8* P[4]-specific (AQTRYAPVNWGHGEINDST+ KYGGRVWTFHGETPRATTDSS+ CNEYINNGLPPIQNTR) or a VP8* P[6]-specific (LDGPYQPTNFKPPND+ NVTNQSRQYTLFGETKQITVENNTNK+HGETPNATTDYSST) synthetic peptide mixture conjugated to keyhole limpet hemocyanin. Serum samples were collected after six vaccinations on day 66, and VP8* genotype-specific total IgG antibodies were affinity purified using VP8* P[8]/VP8* P[4]-specific or VP8* P[6]-specific peptides coupled to SulfoLink columns. All in-life experimental procedures undertaken during the course of the immunization study were conducted in accordance with German laws and guidelines for animal protection and appropriate local and national approvals as well as in accordance with Directive 2010/63/EU of the European Parliament and of the Council of 22 September 2010 on the protection of animals used for scientific purposes.

### *In vitro* protein expression

For expression analysis of mRNA or pDNA encoded proteins in cell culture, HEK 293T or HeLa cells were seeded either in 6-well plates at a density of 3 × 10^5^ cells/well (used for mRNA transfection; Fig. 2b), in T75 flasks at a density of 3 × 10^6^ cells/flask (used for mRNA transfection; Fig. 3c, Supplementary Fig. 3), or in 24-well plates at a density of 1 × 10^5^ cells/well (used for pDNA transfection; Supplementary Fig. 2). 24 hours later, cells were transfected under serum-free conditions either with 2 µg of mRNA per well (6- well plate), with 17 µg of mRNA per flask, or with 1 µg of pDNA per well (24-well plate) via lipofection. For this, mRNAs and pDNAs were complexed with Lipofectamine^TM^ 2000 (Invitrogen) at a ratio of 1:1.5 and 1:2 (w/v), respectively, and transfected into cells according to the manufacturer’s protocol. Protein expression in cell culture supernatants or cell lysates was assessed 48 hours post transfection via SDS- PAGE^70^ and western blotting. Supernatants were filtered through 0.2 µm pore-size PES syringe filters (Sarstedt) and cell pellets were lysed in 1× Laemmli buffer before proteins were separated on 4-20% Mini- PROTEAN^®^ TGX^TM^ Precast Protein Gels (Bio-Rad) and transferred to a nitrocellulose membrane (Odyssey^®^ nitrocellulose membrane, pore-size: 0.22 µm; LI-COR). Specific proteins were detected using rabbit anti-VP8* P[8]/P[4] or P[6] polyclonal antibodies (1:1,000; generated in this study by Aldevron Freiburg GmbH), guinea pig anti-LS-P2-VP8* P[8] polyclonal antiserum (1:500, day 196 serum sample from an animal vaccinated with LS-P2-VP8* P[8] mRNA vaccine; generated in this study by Covance Laboratories Ltd.), and mouse anti-alpha tubulin antibody (1:1,000; Abcam, Cat. ab7291), followed by goat anti-rabbit IgG IRDye^®^ 800CW or 680RD (1:10,000; LI-COR, Cat. 926-32211 or Cat. 926-68071), donkey anti-guinea pig IgG IRDye^®^ 800CW (1:10,000; LI-COR, Cat. 925-32411), and goat anti-mouse IgG IRDye^®^ 680RD (1:10,000; LI-COR, Cat. 926-68070), respectively. Protein detection and image processing were carried out in an Odyssey^®^ CLx Imaging System and LI-COR’s Image Studio^TM^ Lite version 5.2.5 according to the manufacturer’s instructions.

### Density gradient ultracentrifugation

Potential nanoparticles derived from supernatants of HeLa or HEK 293T cell cultures transfected with mRNAs encoding either LS-P2-VP8* P[8] or exclusively full-length lumazine synthase including the two point mutations described above (LS only) were purified by density gradient ultracentrifugation. For this purpose, cell culture supernatants were filtered 48 hours post transfection as described above and cOmplete, Mini, EDTA-free Protease Inhibitor Cocktail (PIC) Tablets were added (1 tablet/10 mL supernatant; Roche). Approximately 2 mL of supernatant were applied onto an isopycnic OptiPrep^TM^-iodixanol (Sigma- Aldrich) density gradient (1.6 mL each of 60%, 50%, 40%, 30%, 20%, and 10% iodixanol in PBS with PIC). After ultracentrifugation (3 hours; 200,000 × g; 4°C; ultracentrifuge: Sorvall WX80; rotor: Sorvall TH-641), seven density gradient fractions (F1 to F7) were collected from the top (F1: lowest iodixanol concentration) to the bottom (F7: highest iodixanol concentration) and stored at −80°C until further analysis.

### Transmission electron microscopy

Potential LS-P2-VP8* P[8] nanoparticles were analyzed by transmission electron microscopy after purification by density gradient ultracentrifugation. For this purpose, LS-P2-VP8* P[8] containing density gradient fractions 3 and 4 (confirmed by SDS PAGE and western blotting) were thawed, pooled and concentrated to ∼0.5 mg/ml in 1× PBS at pH 7.4. Lacey grids with a thin continuous carbon film (Electron Microscopy Sciences, Hatfield, PA), 400 mesh, were negatively glow discharged under vacuum with 20- 25 mA current for 30 s. A 3.0 µL aliquot of sample was applied to these grids at 4°C and 100% humidity, blotted for 3-3.5 s and plunge frozen in liquid ethane using a Vitrobot Mark IV specimen preparation unit (FEI Company, Hillsboro, OR).

Vitrified grids were imaged using an FEI Titan Krios cryo-electron microscope operating at 300 keV equipped with a Gatan K2 summit direct detector and a post-specimen energy filter. Micrographs were collected at ∼130,000× magnification with a pixel size of 0.53Å per pixel in super-resolution mode. Images were collected at a dose rate of ∼8e^−^/pixel/s with 200 ms exposure per frame and 50 frames per image. Estimated defocus ranged from 0.75 to 3 µm. A total of 1291 micrographs was collected using the automated data collection software Leginon^71^.

All data processing steps were performed within Relion (version 3.1) software setup^72^. Frame alignment and dose-weighting were performed using MotionCor2^73^, and CTF estimation was done using CTFFIND4^74^ within the Relion wrapper. A total of 135718 particles were picked using the LOG-based automatic particle picking routine. Particles were initially extracted at 16× binning with a pixel size of 8.48 Å/pixel. Unsupervised 2D classification of the binned particle stack was carried out. Classes that showed clearly defined circular and likely icosahedral featured particles were selected. A total of 3622 particles were selected and subjected to a second round of 2D classification. From this round, 1936 particles were selected. The previously published cryo-EM structure of lumazine synthase icosahedral nanoparticle (EMD-3538) was used as initial model. The initial model was low-pass filtered to 60 Å and refinement was performed on the selected particle stack with icosahedral symmetry imposed. Sequential unbinning of particle stacks along with refinement was carried out till bin2. Particle polishing and per-micrograph ctf refinement was then carried out to improve the 3D density map. Subsequent map sharpening and post processing in Relion resulted in a 2.6 Å structure using the ‘gold-standard’ FSC cutoff of 0.143.

The atomic model of *Aquifex aeolicus* lumazine synthase nanoparticle (PDB ID: 5MPP)^43^ was rigid body fitted into the LS-P2-VP8* P[8] 3D map using UCSF Chimera^75^. The atomic model agreed well with the map. Analysis of the map also revealed that there was no density visible for the P2-VP8* portion. The cryo-EM map was further sharpened using the Autosharpen program^76^ in the Phenix software suite^77^ in an attempt to enhance map features. The rigid bidy-fitted lumazine synthase PDB model was then refined using Realspace Refinement in the Phenix package using default restraints but without any annealing or ADP refinement^77^. The cryo-EM map features are fully occupied by the lumazine synthase model with no extra density visible for P2-VP8* suggesting that the P2-VP8* portion averaged out during the reconstruction process likely due to its flexible linkage to the lumazine synthase protein. The data collection, refinement parameters and final model statistics are summarized in Supplementary Table 1.

### Statistical analyses

All statistical analyses were performed using GraphPad Prism version 9.4.1 for Windows (GraphPad Software). The statistical significance of differences between groups was examined using an ordinary one- way ANOVA followed either by Tukey’s multiple comparison post test when all groups were compared with each other or by Šídák’s multiple comparison post test when pre-selected pairwise analysis was used. All binding and neutralizing antibody data were log_10_-transformed for analysis. To test for normal or lognormal distribution, D’Agostino & Pearson, Anderson-Darling, Shapiro-Wilk, and/or Kolmogorov-Smirnov tests were used. Each *P* value was multiplicity adjusted to account for multiple comparisons. Differences were considered significant at adjusted *P* values of <0.05. Supplementary Table 2 provides a detailed summary of ANOVA results including multiple comparison testing for each statistical analysis.

## DATA AVAILABILITY

The authors declare that all relevant data supporting the findings of this study are available within the paper and its supplementary information files. Additional information and underlying data are available from the corresponding author upon reasonable request. The cryo-EM data and structure of the LS-P2-VP8* P[8] nanoparticle generated in this study have been submitted to the Electron Microscopy Data Bank (EMDB) under the accession code EMD-28807 and to the Protein Data Bank (PDB) under the accession code 8F25. These are publicly available as of the date of publication.

## ACKNOWLEDGEMENTS

Funding for this study was provided by the Bill & Melinda Gates Foundation (INV-020846 and OPP- 1126258). We thank Holger Kanzler, Carl Kirkwood and Amy Weiner (Bill & Melinda Gates Foundation) for their expert advice and excellent support throughout the project. Thanks to Stanley Cryz, Jessica White, Kyle Lakatos and Marcus Estrada (PATH) for providing the P2-VP8* proteins and guidance on alum- adjuvantation. We acknowledge Nicole Meyer (Cincinnati Children’s Hospital Medical Center) for her assistance in performing the rotavirus neutralization assays. Thanks to Jennifer Oduro and Jonas Füner (Preclinics) as well as Isam Sharif (Covance; now Labcorp) for their outstanding commitment and expertise in conducting the *in vivo* studies. A very special thanks goes to Julia Schröder, Ariane Heiler, Jessica Lahm and Julia Führer (CureVac) for performing most of the experiments and their relentless and expert support in the lab. An additional special thanks to Ariane Heiler for her enthusiasm and expertise in performing mutational analyses. Thanks to Andreas Theß, Moritz Thran and Wolfgang Große (CureVac) for their support with generating mRNA constructs, Susanne Braeuer and Aniela Wochner (CureVac) for mRNA production, Alba Martinez Munuera, Annachiara Greco and Marius Busche (CureVac) for all their excellent work on project planning and communication, and thanks to Igor Splawski (CureVac) for critically reading the manuscript.

## AUTHOR CONTRIBUTIONS

S.Ro., B.P., and S.Ra. conceived and conceptualized the work and strategy. S.Ra designed the mRNA vaccines. S.Ro. and S.Ra. designed the *in vitro* and *in vivo* studies and analyzed and interpreted the data. V.M.P. and K.K.L. designed, conducted, and analyzed the electron microscopy experiments. M.M.M. performed and analyzed the rotavirus neutralization assays. S.Ro. and S.Ra. wrote the manuscript with input from all authors. All authors reviewed the manuscript and approved the final version.

## COMPETING INTERESTS

S.Ro., B.P., and S.Ra. are employees of CureVac SE, Tübingen, Germany, a publicly listed company developing mRNA-based vaccines and immunotherapeutics, and are inventors on several patents on mRNA vaccination and use thereof. S.Ro. and B.P. hold shares in the company. The other authors declare no competing interests.

## SUPPLEMENTARY INFORMATION

**Supplementary Fig. 1.**
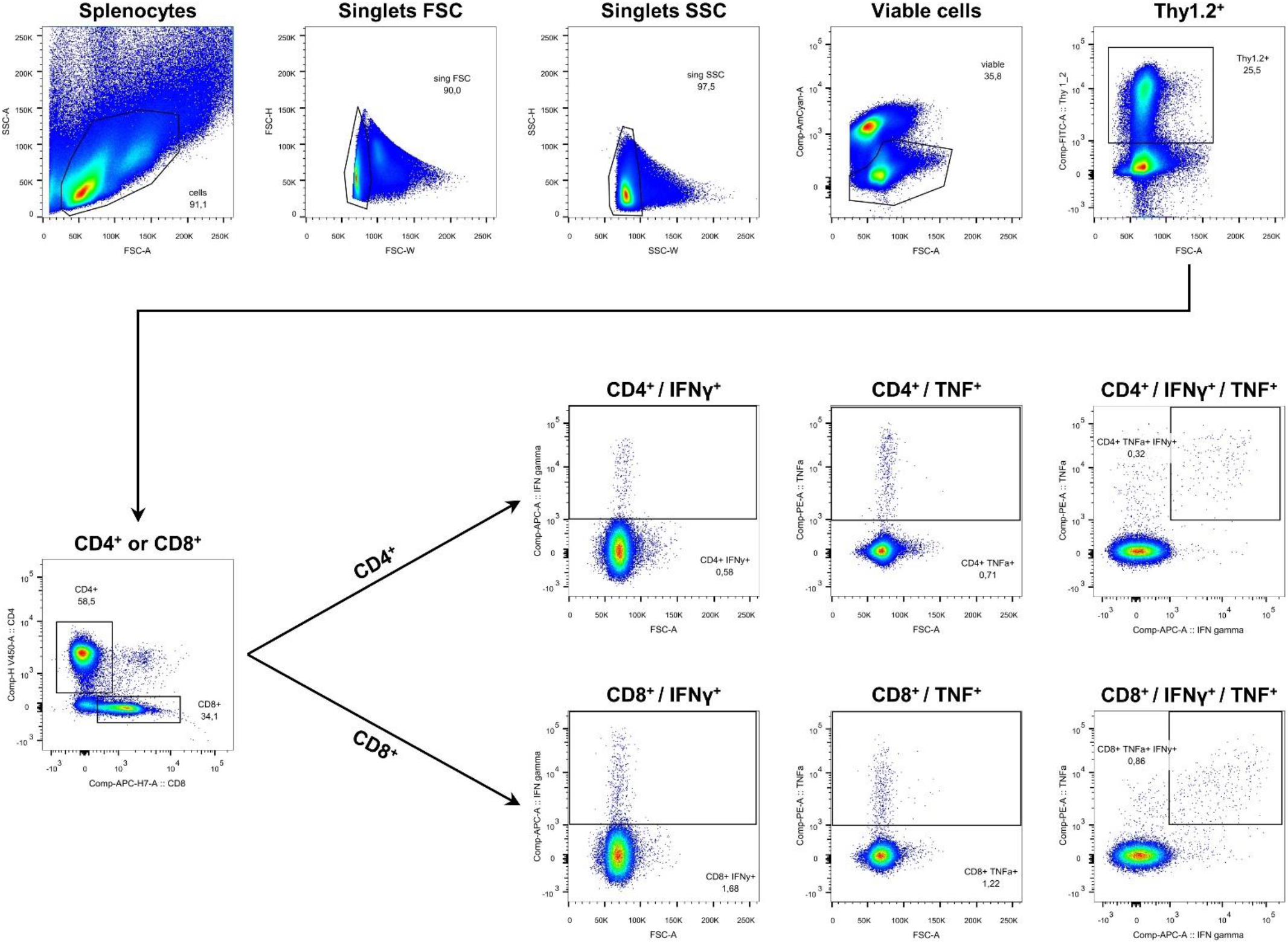
Gating strategy for T cell analysis. Female BALB/c mice were vaccinated and splenocytes were isolated as described in Fig. 1. Multifunctional IFN-γ/TNF-positive CD4^+^ and CD8^+^ T cells were analyzed in splenocytes stimulated with VP8*-specific peptides followed by intracellular cytokine staining and detection by flow cytometry. The flow blots shown illustrate the gating strategy using splenocytes isolated from a mouse vaccinated with P2-VP8* P[8] mRNA vaccine as an example. T cells were characterized as singlets, viable cells, Thy1.2^+^, CD4^+^ or CD8^+^ and subdivided into multifunctional T cell subsets based on their expression of IFN-γ and TNF.

**Supplementary Fig. 2.**
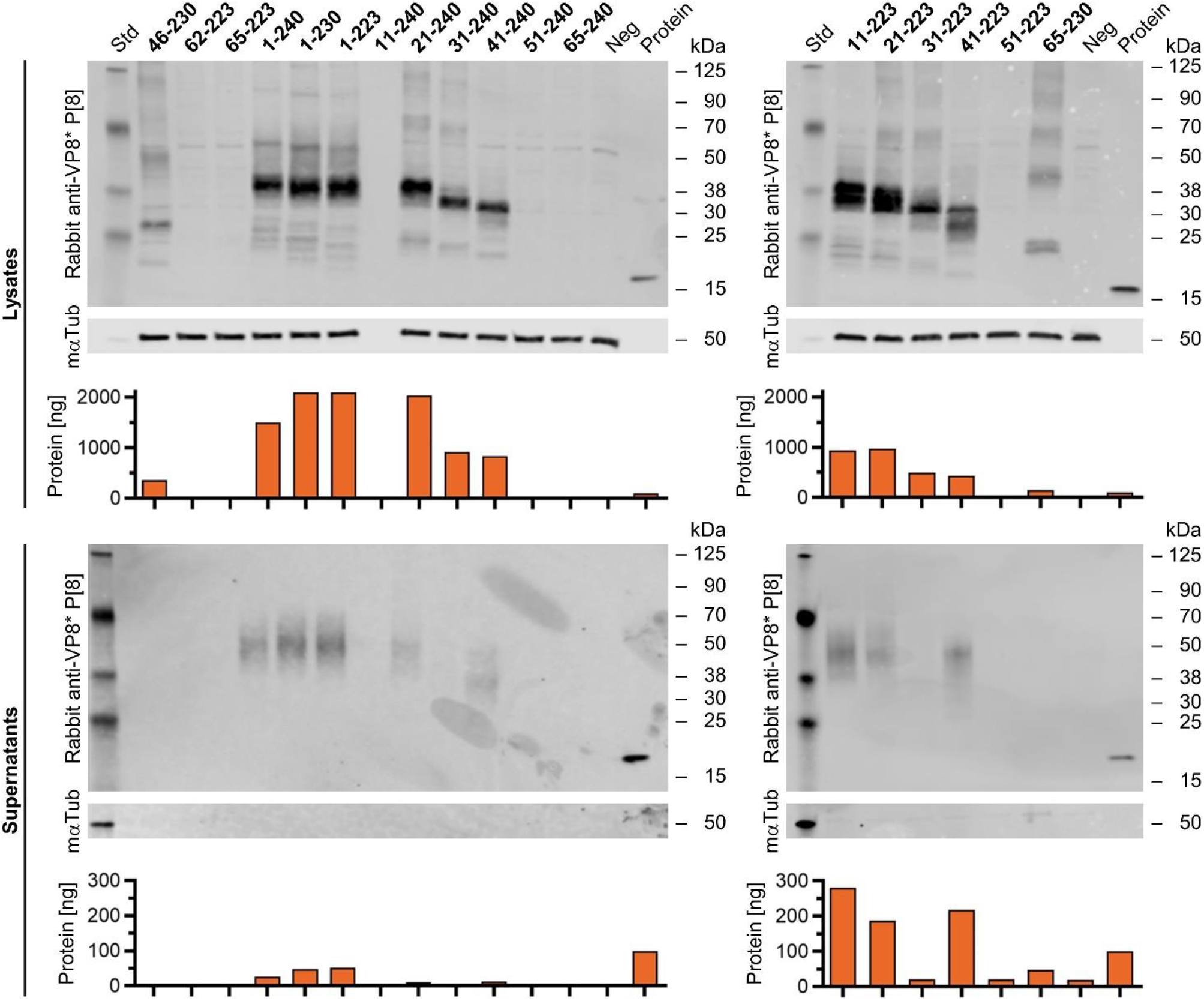
Expression and secretion levels of proteins translated from pDNAs encoding for different amino acid lengths of VP8* P[8] differed considerably. VP8* expression in HEK 293T cells transfected with different pDNA constructs as indicated was analyzed from cell lysates or supernatants via western blotting 48 hours post transfection. A supernatant or cell lysate from HEK 293T cells transfected with water served as negative control (Neg) and 100 ng of P2-VP8* P[8] protein loaded to the gel was used as positive control (Protein). Tubulin was employed as loading control. Protein expression was quantified using the Image Studio^TM^ Lite version 5.2.5 software and normalized to the signal corresponding to 100 ng of P2-VP8* P[8] protein. mαTub: mouse anti-alpha tubulin antibody.

**Supplementary Fig. 3.**
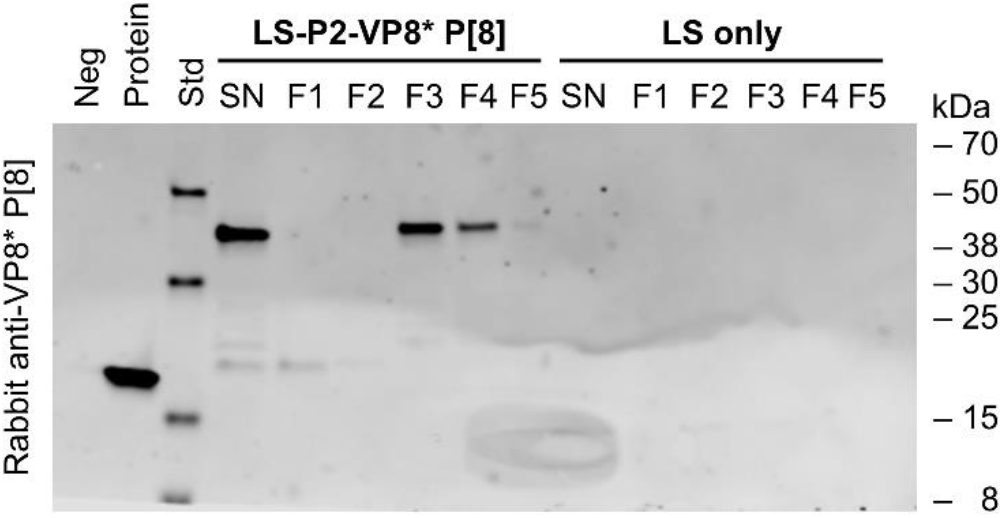
VP8*-specific signals were only detectable in cellular supernatants transfected with LS-P2-VP8* P[8] mRNA, both before and after ultracentrifugation (compare Fig. 2c). Cell supernatants of HeLa cells transfected with LS-P2-VP8* P[8] or LS only mRNA were subjected to density gradient ultracentrifugation using an isopycnic iodixanol density gradient 48 hours post transfection. After ultracentrifugation, equivalent volumes of five density gradient fractions with increasing iodixanol concentrations (F1 to F5) or unpurified supernatants (SN) were analyzed for VP8* expression via western blotting. A supernatant from HeLa cells transfected with water served as negative control (Neg) and 100 ng of P2-VP8* P[8] protein loaded to the gel was used as positive control (Protein). LS: lumazine synthase.

**Supplementary Fig. 4.**
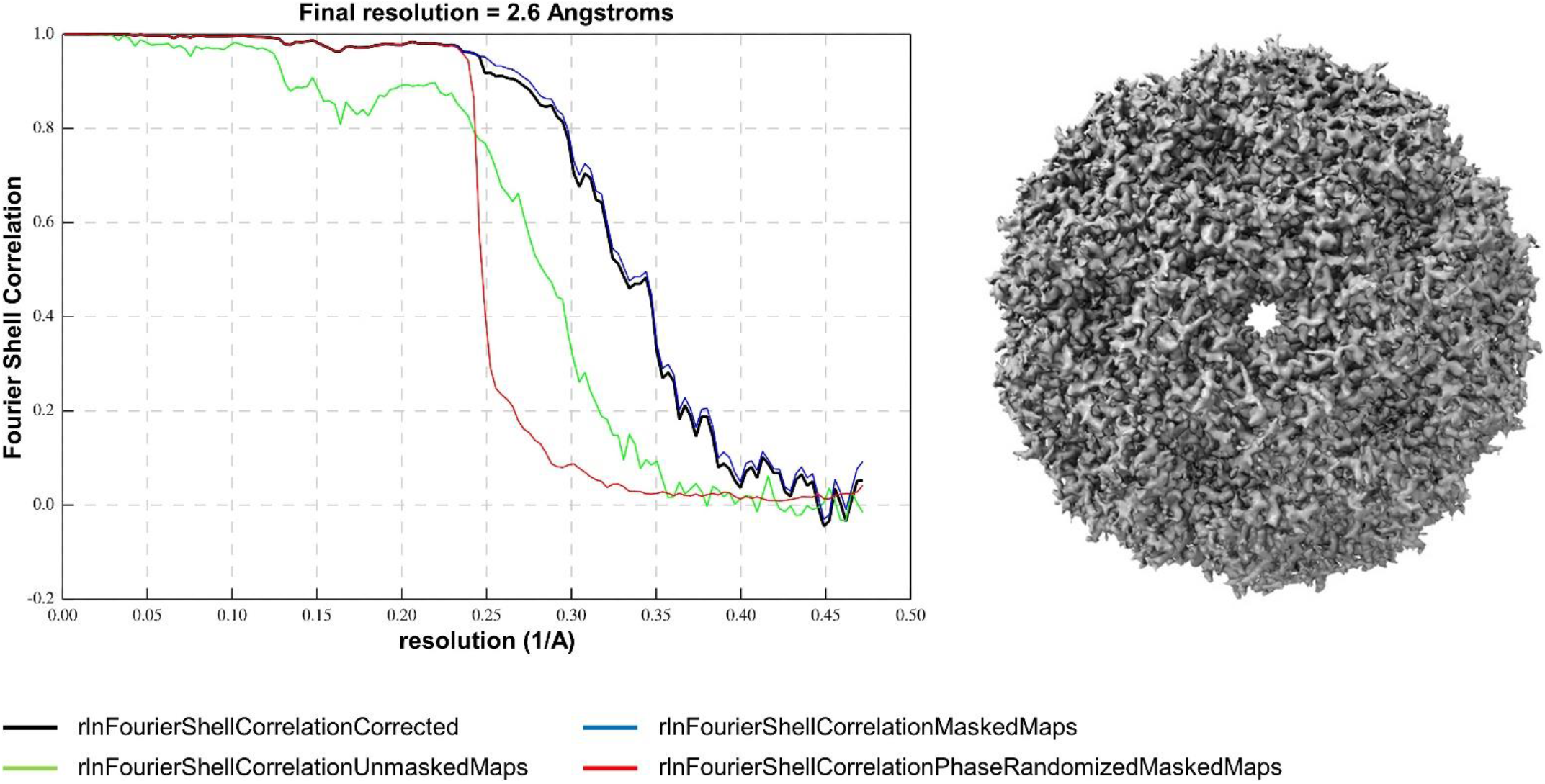
Resolution estimation for LS-P2-VP8* P[8] cryo-EM structure. Fourier Shell Correlation (FSC) curve for the LS-P2-VP8* P[8] nanoparticle cryo-EM structure (left) along with a 2D snapshot of density map (right).

**Supplementary Fig. 5.**
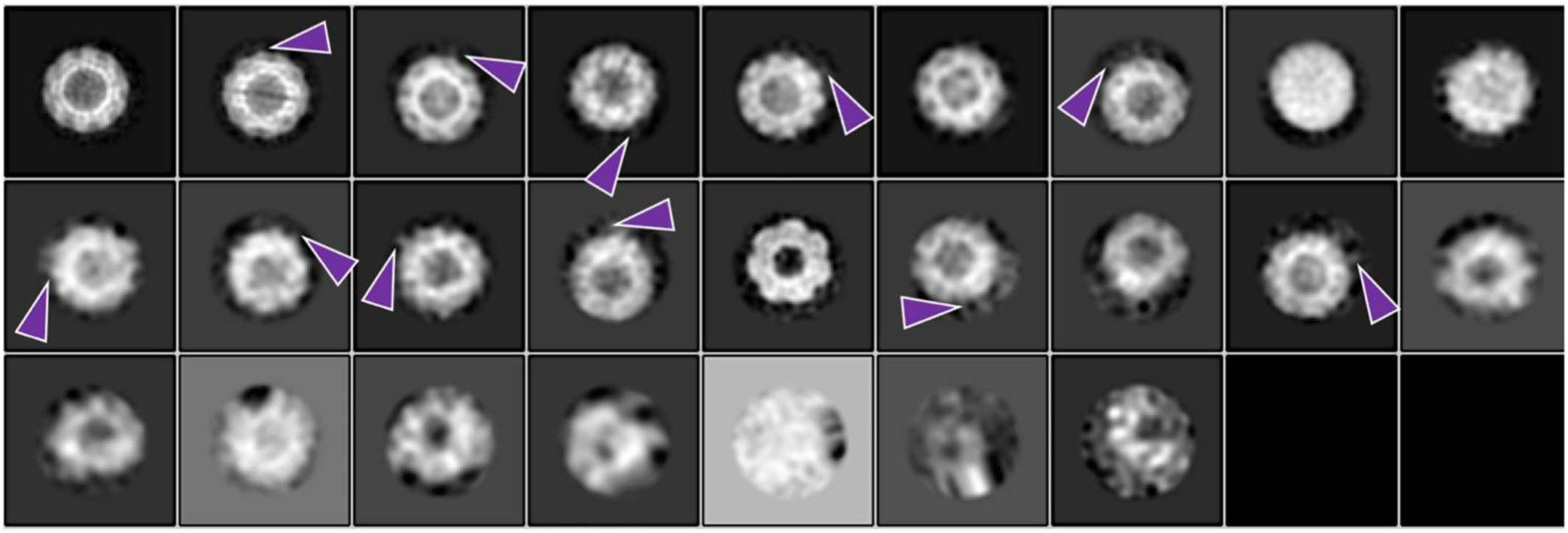
2D class averages from LS-P2-VP8* P[8] cryo-EM data demonstrated nonuniform density protruding from nanoparticle surface. Images of averaged classes obtained from unsupervised 2D classification of LS-P2-VP8* P[8] particle images. Purple arrows point to external weak densities seen protruding from the surface of the nanoparticles. White is high density in all the panels.

**Supplementary Fig. 6.**
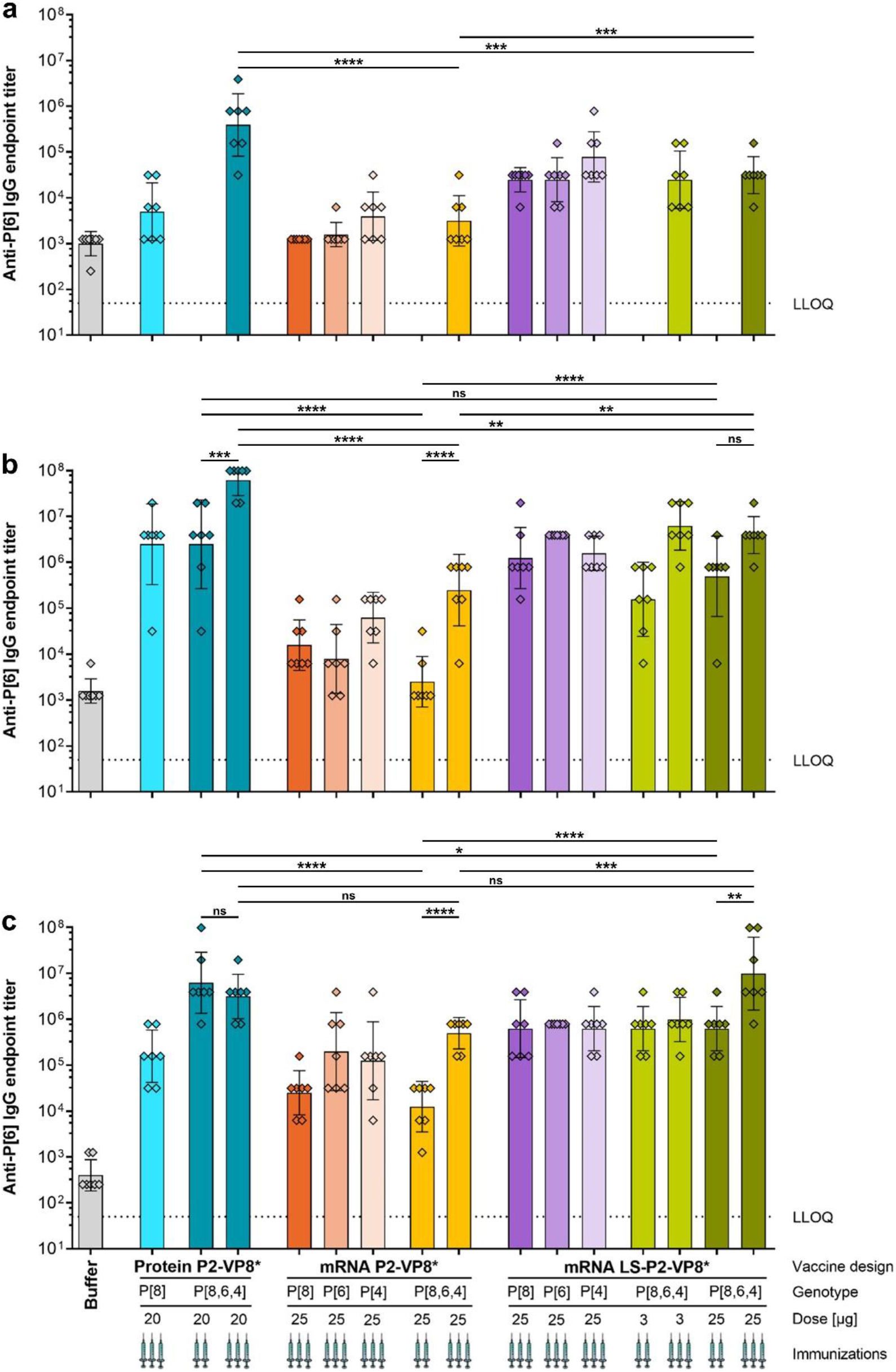
Trivalent LS-P2-VP8* mRNA vaccine encoding genotypes P[8], P[6] and P[4] of VP8* elicited high titers of binding antibodies against the P2-VP8* P[6] protein in guinea pigs. Female Dunkin-Hartley guinea pigs (n = 7/group) were vaccinated IM either three times on day 0, day 21, and day 42 or twice on day 21 and day 42 with different doses of monovalent and trivalent alum-adjuvanted P2-VP8* protein vaccines or monovalent and trivalent P2-VP8* or LS-P2-VP8* mRNA vaccines as indicated. The genotype(s) of the encoded VP8* employed for each group, the doses used as well as the number of immunizations administered (indicated by the number of syringe symbols) are displayed beneath each group. Trivalent vaccines are labeled P[8,6,4]. Buffer (0.9% NaCl) vaccinated animals served as negative controls. P2-VP8* P[6]-specific binding antibodies are displayed as endpoint titers for total IgG in serum collected on day 21 **a**, day 42 **b**, or day 70 **c**. Each square represents an individual animal and bars depict geometric means with geometric SD. Dotted lines indicate the lower limit of quantification (LLOQ). Individual time points were statistically analyzed using an ordinary one-way ANOVA followed by Šídák’s multiple comparison post test. Significant differences between groups are marked by asterisks (\**P*<0.05, ** *P*<0.01, *** *P*<0.001, **** *P*<0.0001; ns: not significant).

**Supplementary Fig. 7.**
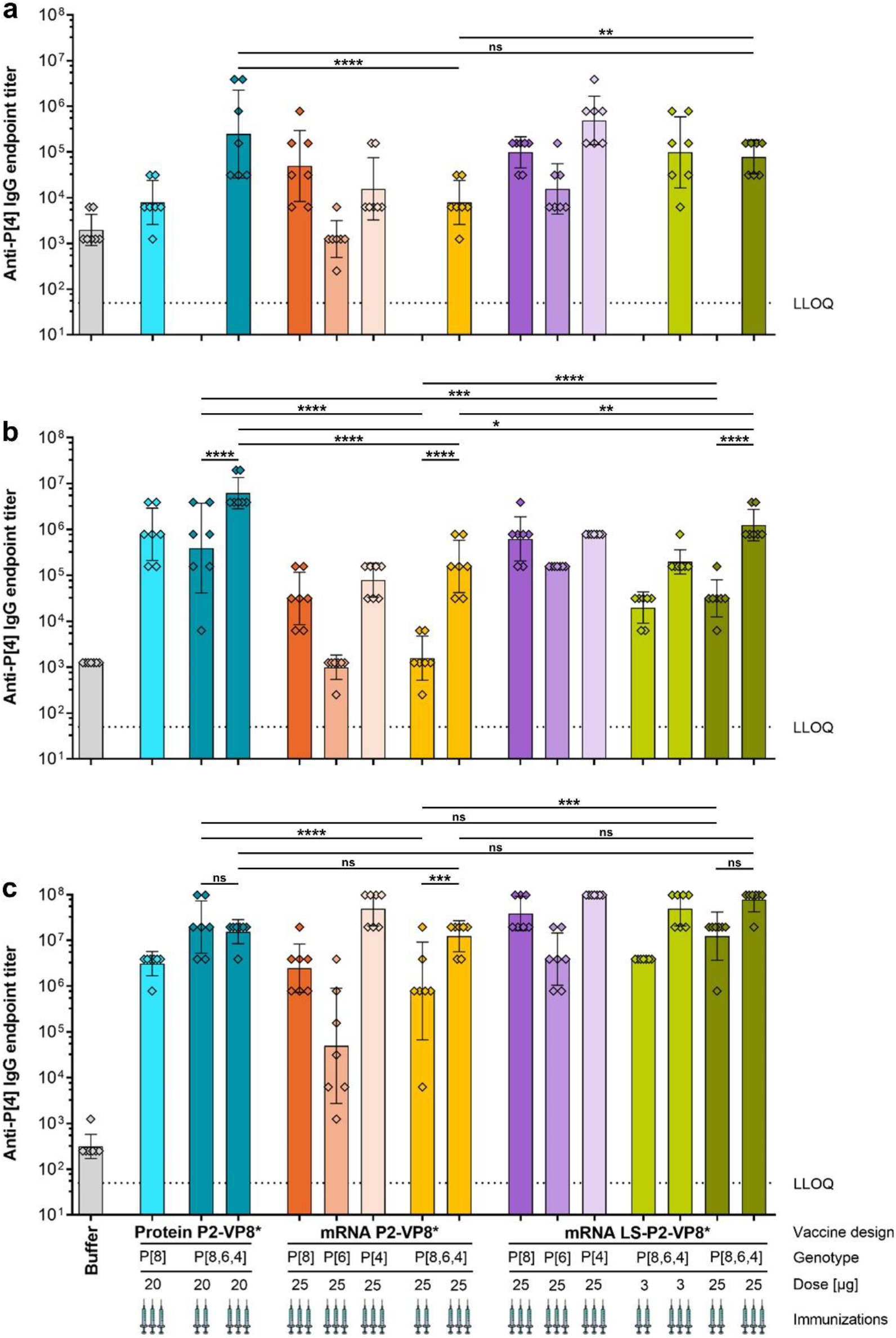
Trivalent LS-P2-VP8* mRNA vaccine encoding genotypes P[8], P[6] and P[4] of VP8* elicited high titers of binding antibodies against the P2-VP8* P[4] protein in guinea pigs. Female Dunkin-Hartley guinea pigs (n = 7/group) were vaccinated IM either three times on day 0, day 21, and day 42 or twice on day 21 and day 42 with different doses of monovalent and trivalent alum-adjuvanted P2-VP8* protein vaccines or monovalent and trivalent P2-VP8* or LS-P2-VP8* mRNA vaccines as indicated. The genotype(s) of the encoded VP8* employed for each group, the doses used as well as the number of immunizations administered (indicated by the number of syringe symbols) are displayed beneath each group. Trivalent vaccines are labeled P[8,6,4]. Buffer (0.9% NaCl) vaccinated animals served as negative controls. P2-VP8* P[4]-specific binding antibodies are displayed as endpoint titers for total IgG in serum collected on day 21 **a**, day 42 **b**, or day 70 **c**. Each square represents an individual animal and bars depict geometric means with geometric SD. Dotted lines indicate the lower limit of quantification (LLOQ). Individual time points were statistically analyzed using an ordinary one-way ANOVA followed by Šídák’s multiple comparison post test. Significant differences between groups are marked by asterisks (\**P*<0.05, ** *P*<0.01, *** *P*<0.001, **** *P*<0.0001; ns: not significant).

**Supplementary Table 1.** Cryo-EM data collection, refinement and validation statistics.

|  | LS-P2-VP8* P[8] nanoparticles<br>(EMDB-28807), (PDB 8F25) |
| --- | --- |
| **Data collection and processing** |  |
| Magnification | 130000× |
| Voltage (kV) | 300 kV |
| Electron exposure (e<sup>-</sup>/Å<sup>2</sup>) | 284.8 |
| Defocus range (μm) | -0.75-3 |
| Pixel size (Å) | 0.53 |
| Symmetry imposed | Icosahedral |
| Initial particle images (no.) | 135718 |
| Final particle images (no.) | 1936 |
| Map resolution (Å) | 2.61 (0.143) |
| FSC threshold |  |
| Map resolution range (Å) | - |
| **Refinement** |  |
| Initial model used (PDB code) | 5MPP |
| Model resolution (Å) | 2.6 |
| FSC threshold |  |
| Model resolution range (Å) | - |
| Map sharpening *B* factor (Å<sup>2</sup>) | -81.9253 |
| R.m.s. deviations |  |
| Bond lengths (Å) | 0.007 |
| Bond angles (°) | 0.685 |
| Validation |  |
| MolProbity score | 4.04 |
| Clashscore | 0.0 |
| Poor rotamers (%) |  |
| Ramachandran plot |  |
| Favored (%) | 96.04 |
| Allowed (%) | 3.96 |
| Disallowed (%) | 0 |

**Supplementary Table 2 | Comprehensive information on the statistical analyses.** The accompanying Excel file provides a detailed summary of ANOVA results including multiple comparison testing for each statistical analysis performed in this study.

## Notes

### Competing Interest Statement

Funding for this study was provided by the Bill & Melinda Gates Foundation (INV-020846 and OPP-1126258). S.Ro., B.P., and S.Ra. are employees of CureVac SE, Tuebingen, Germany, a publicly listed company developing mRNA-based vaccines and immunotherapeutics, and are inventors on several patents on mRNA vaccination and use thereof. S.Ro. and B.P. hold shares in the company. The other authors declare no competing interests.

